# Composition structures and biologically meaningful logics: plausibility and relevance in bipartite models of gene regulation

**DOI:** 10.1101/2022.05.07.491027

**Authors:** Yasharth Yadav, Ajay Subbaroyan, Olivier C. Martin, Areejit Samal

**Author notes:** Y.Y. and Aj.S. contributed equally to this work and should be considered as Joint-First authors.

## Abstract

Boolean network models have widely been used to study the dynamics of gene regulatory networks. However, such models are coarse-grained to an extent that they abstract away molecular specificities of gene regulation. In contrast, *bipartite* Boolean network models of gene regulation explicitly distinguish genes from transcription factors (TFs). In such models, multiple TFs may simultaneously contribute to the regulation of a gene by forming heteromeric complexes. The formation of heteromeric complexes gives rise to *composition structures* in the corresponding bipartite network. Remarkably, composition structures can severely restrict the number of Boolean functions (BFs) that can be assigned to a gene. The introduction of bipartite Boolean network models is relatively recent, and so far an empirical investigation of their biological plausibility is lacking. Here, we estimate the prevalence of composition structures arising through heteromeric complexes in *Homo sapiens*. Moreover, we present an additional mechanism by which composition structures arise as a result of multiple TFs binding to the *cis*-regulatory regions of a gene and we provide empirical support for this mechanism. Next, we compare the restriction in BFs imposed by composition structures and by biologically meaningful properties. We find that two types of minimally complex BFs, namely nested canalyzing functions (NCFs) and read-once functions (RoFs), are more restrictive than composition structures. Finally, using a compiled dataset of 2687 BFs from published models, we find that composition structures are highly enriched in real biological networks, but that this enrichment is most likely driven by NCFs and RoFs.

## I. INTRODUCTION

Transcriptional regulation is one of the most fundamental mechanisms for the control of gene expression in eukaryotes [1]. This realization has led to the construction and analysis of the structure and dynamics of transcriptional gene regulatory networks [2–5]. One of the most widely used frameworks for studying the dynamics of such gene regulatory networks are Boolean networks. Stuart Kauffman [6, 7] and René Thomas [8, 9] pioneered the use of Boolean models to reproduce some of the essential dynamical behaviours of living systems such as fixed point dynamics and cyclic dynamics [4, 10, 11]. Over time Boolean modeling has gained a wide appeal and has thus been extended to capture the dynamics of other types of biological networks such as signalling and metabolic networks [12–14].

Until the turn of this century, the paucity of empirical data on the structure (including combinatorial regulation) of real biological networks mandated a statistical approach based on ensembles of random Boolean networks for attempting to infer the dynamics of gene regulatory networks [6, 10, 15–17]. Random Boolean networks are typically defined by placing interactions (directed edges) between randomly chosen genes (nodes) and assigning random logical update rules to those nodes. However, mounting evidence over the past two decades, obtained via biological network reconstruction using large-scale data from high-throughput experiments [3, 18], has shown that the architecture of real gene networks is far from random, both for their network structure [3, 5, 13, 18–23] and for their logical update rules [11, 13, 24-31], i.e., the Boolean functions (BFs) assigned to each associated gene.

Further investigation into the nature of Boolean functions capturing gene regulation revealed that regulatory Boolean logics possess certain properties based on which they are naturally classified into the so-called biologically meaningful types of BFs: unate functions (UFs) [32], canalyzing functions (CFs) [33] and nested canalyzing functions (NCFs) [10, 34, 35]. Stuart Kauffman, in his book titled “The Origin of Life” [33], propounded the idea that canalyzing functions reflect the “chemical simplicity” of the underlying molecular regulatory mechanisms in gene networks. Using Kauffman’s proposition as a premise, Subbaroyan *et al.* [23] showed that NCFs and RoFs (Read-once functions) minimize two notions of complexity of Boolean functions, namely, the average sensitivity and the Boolean complexity, respectively. This effort has led to the addition of a new type of BF previously unexplored in Boolean models of living systems, the Read-once function (RoF), to the list of biologically meaningful BFs. Remarkably, the ‘simplest’ logics, NCFs and RoFs also form the most restrictive subset of BFs among other biologically meaningful BFs in the space of all BFs [23].

The Boolean network model of gene regulation is surely a coarse-grained picture of biological reality. Attempts to incorporate more realistic features within the Boolean framework [36, 37] have been proposed. One such effort has been to use *bipartite* Boolean network models to explicitly capture two types of molecular species, namely, genes and transcription factors involved in transcriptional regulation [37–40]. In particular, Hannam *et al.* [37] introduced a bipartite Boolean network model with genes and TFs, wherein the state of a gene is determined by a BF of the states of its input TFs while the (on or off) state of a TF is in turn determined by a BF of the states of the genes directly contributing to its synthesis. Fink and Hannam [41] capture this gene-TF-gene interaction via subgraphs called “composition structures” in the bipartite Boolean network and elucidate how such a composition structure allows for a ‘composition’ of BFs to be defined on genes. Interestingly, composition structures severely restrict the number of allowed logic rules in the space of all Boolean functions [41]. The restrictive nature of the composition of BFs, albeit in unipartite Boolean models, has also been previously explored by Shmulevich *et al.* [42].

Currently, the bipartite Boolean models proposed [37, 40, 41] for transcriptional gene regulation can at best be considered as only theoretical propositions without a solid grounding in empirical evidence. Our work approaches the question of prevalence of composition structures in real gene regulatory networks from a data-centric perspective. The central theme of our work is thus to examine how plausible it is for both composition structures and composed BFs to occur in real networks by analyzing associated data. We begin by estimating the prevalence of composition structures arising in two different scenarios of gene regulation. The first scenario is gene regulation by heteromeric protein complexes which act as transcription factors [37, 40]. The other scenario, which is a novel aspect of this work, is accounting for transcriptional regulation via *cis*-regulatory elements, in particular, promoters and enhancers [43, 44]), that can be bound by transcription factors.

Next, we build upon the work of Fink and Hannam [41] on Boolean compositions and augment their approach for counting the number of possible BFs under Boolean compositions by accounting for the fact that the different input variables are distinguishable and so are nonequivalent under permutation. We then compare the restriction in the logical rules in gene regulatory networks due to Boolean compositions with the restriction due to different types of biologically meaningful BFs, and thereafter analyze how often Boolean compositions display biologically meaningful properties. Finally, we evaluate the enrichment (depletion) and *relative* enrichment (depletion) of composed BFs in a compiled empirical dataset of 2687 BFs from reconstructed Boolean models of biological systems. Overall, our analyses show that bipartite Boolean models are more flexible than envisaged previously as they can accommodate more plausible modes of gene regulation such as via *cis*-regulatory elements, and that biologically meaningful BFs, in particular NCFs, impose relatively stricter restrictions on the space of allowed regulatory logic in comparison to Boolean composition structures.

## II. THEORETICAL BACKGROUND

### A. Boolean models of gene regulatory networks

A Boolean model of a gene regulatory network consists of nodes (or vertices) and directed links (or edges) wherein the nodes correspond to genes and a directed link towards any gene captures the regulation of that output gene by a input gene from which the link arises [6–8, 33]. In a Boolean network model, the allowed states for any node are analogous to that of a switch which can be either ‘on’ or ‘off’, and therefore, the state of a node is given by a Boolean variable *x* that can take the values 1 or 0, respectively. The dynamics of any Boolean network model is determined by two factors, namely: *update rules* or *Boolean functions* (BFs) which are assigned to each node, and the update scheme employed (*synchronous* [33] or *asynchronous* [45]). The state of a node *j* in the network with *k* inputs at time *t* + 1 is given by a BF *f* = *f_j_*(*x*_1_, *x*_2_,…, *x_k_*) having *k* input variables, where *x_i_* ∈ {0, 1} are the states of each of the *k* inputs at time *t*. This BF maps the 2^*k*^ different possibilities for the *k* input variables to output values 0 or 1, i.e., *f*: {0, 1}^*k*^ → {0, 1}. Thus, the number of possible BFs that can be assigned to a node with *k* inputs is 2^2^*k*^^.

A BF *f* of *k* inputs can be represented as an algebraic expression consisting of the *k* input variables, combined using the logical operators AND (e.g. *x*_1_ · *x*_2_), OR (e.g. *x*_1_ + *x*_2_) and NOT (e.g. 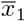). Alternatively, *f* can be represented as a truth table with 2^*k*^ rows, wherein each row corresponds to a possible choice of the set of *k* input variables (see Figure 1). The last entry of each row in the truth table gives the output value for the corresponding realisation of the input variables. Thus, a BF can also be expressed as a binary vector of size 2^*k*^, where each element of the vector corresponds to the output value of the corresponding row of the truth table.

**FIG. 1.**
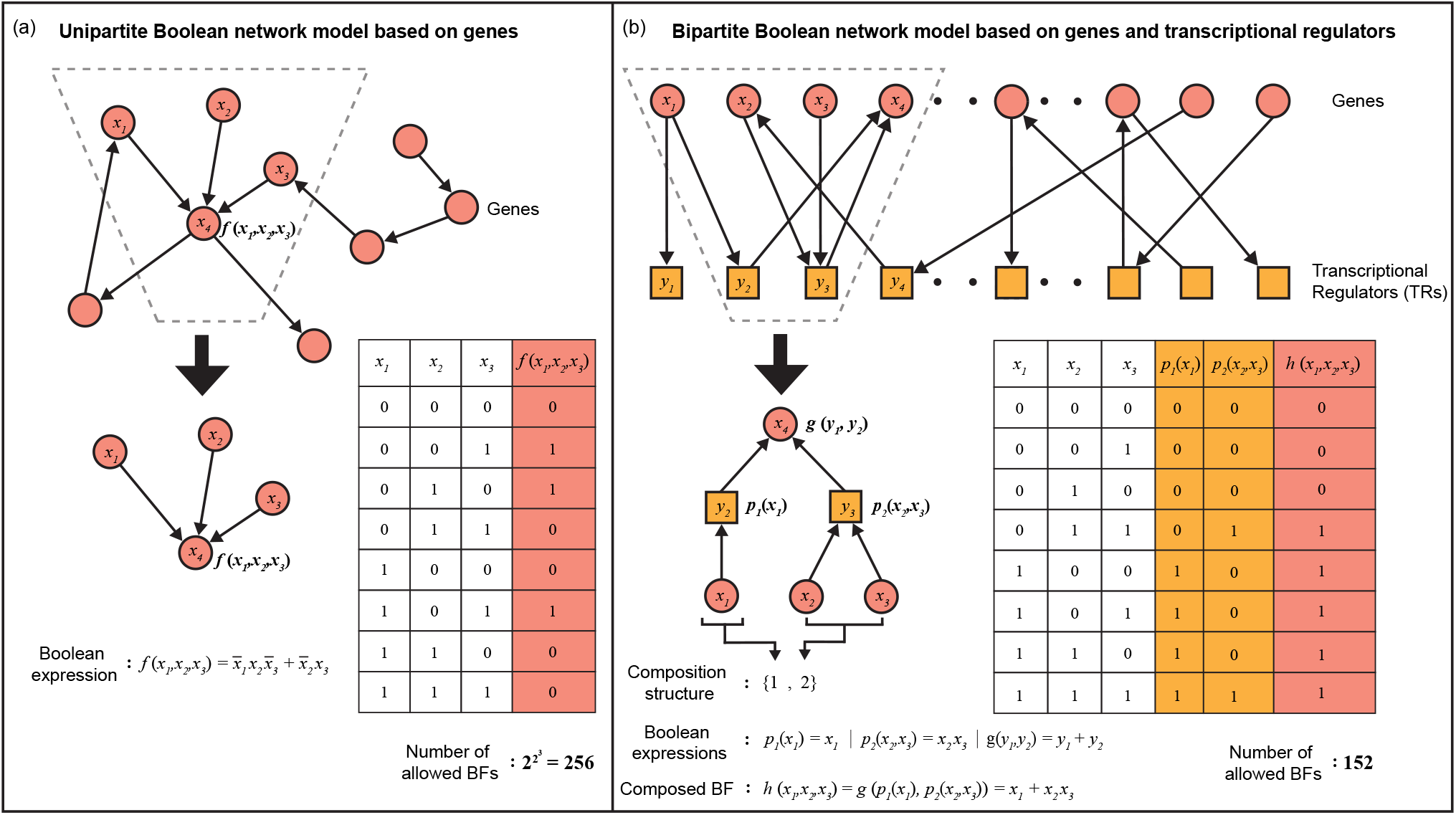
Boolean functions in unipartite versus bipartite network models of transcriptional gene regulation. (a) A unipartite Boolean network model consisting of only genes. The dashed trapezium highlights the subgraph wherein 3 genes with expression states *x*_1_, *x*_2_ and *x*_3_, directly regulate the gene with expression state *x*_4_. Thus, the BF *f* determining the state *x*_4_ of the output gene depends on the states of the 3 input genes *x*_1_, *x*_2_ and *x*_3_, and the truth table for this 3-input BF *f* is shown in the figure. Note that any one of the 2^2^3^^ = 256 possible 3-input BFs can be assigned to BF *f*. (b) A bipartite Boolean network model accounting for the two types of molecular species involved in transcriptional regulation namely, the genes and transcriptional regulators (TRs). In this bipartite Boolean network model, the states of genes are denoted by variables *x*_1_, *x*_2_,…,*x_i_* and the states of TRs are denoted by variables *y*_1_, *y*_2_,…, *y_j_*. The dashed trapezium highlights the subgraph wherein the gene with state *x*_1_ determines the TR with state *y*_1_ according to a 1-input BF *p*_1_(*x*_1_) = *x*_1_, and the genes with states *x*_2_ and *x*_3_ determine the state of the TR *y*_2_ according to a 2-input BF *p*_2_(*x*_2_, *x*_3_) = *x*_2_ *x*_3_. Moreover, the TRs with states *y*_1_ and *y*_2_ in this subgraph directly regulate the gene with state *x*_4_ according to a 2-input BF *g*(*y*_1_, *y*_2_) = *y*_1_ + *y*_2_. Ultimately, the regulation of the output gene with state *x*_4_ depends on the states of the input genes *x*_1_, *x*_2_ and *x*_3_ according to a 3-input BF *h*(*x*_1_, *x*_2_, *x*_3_) = *g*(*p*_1_(*x*_1_),*p*_2_(*x*_2_, *x*_3_)) = *x*_1_ + *x*_2_ *x*_3_. Fink and Hannam [41] called such a subgraph a “composition structure” and the BF *h* corresponding to the subgraph a “composed BF”. The truth table of a composed BF *h* allowed by the composition structure {1, 2} is shown in the figure. Of the 2^2^3^^ = 256 possible 3-input BFs, only 152 BFs can be realised within the composition structure {1, 2}.

### B. Useful properties of Boolean functions

Some properties associated with BFs such as bias or parity, complementarity or being isomorphic [46], may have utility in the biological context [23]. The bias of a BF *f* is the number of ones which occur in its output vector. A BF with an odd (even) bias is said to possess an odd (even) parity. The complement 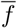 of a BF *f* is obtained by inverting each element in its output vector [46]. For instance, if BF *f* = [0, 0, 1, 0], then its complement 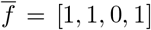. The isomorphisms [46] of a BF *f* are the BFs obtained by permuting and possibly negating the inputs of *f*. For instance, the 4 isomorphisms of the BF with expression *a* + *b* are *a* + *b* itself, *a* + *b, a* + *b*, and *a* + *b*.

### C. Biologically meaningful types of Boolean functions

#### Unate function (UF)

If a BF is monotonically increasing or decreasing for each input *i*, it is said to be a unate function (UF) [32]. Formally, if a BF is monotonically increasing in *x_i_*, then:

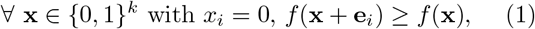

or, if it is monotonically decreasing, then:

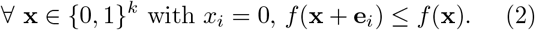

Here **e**_*i*_ ∈ {0, 1}^*k*^ denotes the unit vector having entry 1 for input *i* and 0 for all others.

#### Canalyzing function (CF)

If in a BF there exists at least one input *i* (or input variable *x_i_*) which, when fixed to 0 or 1, fixes the output value, then that BF is said to be a canalyzing function (CF) [33]. Mathematically,

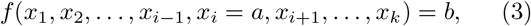

independent of *x_j_* for *j* ≠ *i*. Here, *x_i_* is the canalyzing input variable, *a* is the canalyzing input value, and *b* is the canalyzed output value.

#### Nested canalyzing function (NCF)

A *k*-input BF is said to be a nested canalyzing function (NCF) [10, 47] with respect to some permutation *σ* on its inputs if:

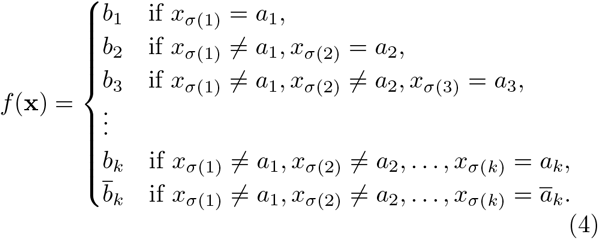

Here, *a*_1_, *a*_2_,…, *a_k_* are the canalyzing input values and *b*_1_, *b*_2_,…, *b_k_* are the canalyzed output values. 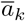 and 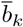 denote the complements of the Boolean values *a_k_* and *b_k_*, respectively.

#### Read-once function (RoF)

If a *k*-input BF *f* can be expressed only using the operators AND, OR and NOT in such a manner that each variable appears exactly once in the Boolean expression, then the BF is said to be a Read-once function (RoF) [23, 48]. Mathematically, for *f* there exists a permutation *σ* on {1, 2,…,*k*} such that after stripping all the parentheses in the Boolean expression for *f* (**x**), we are left with an expression of the form:

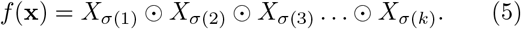

Here, 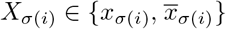 and ⊙ ∈ {∧ (AND), ∨ (OR)}.

### D. Bipartite Boolean network model of transcriptional gene regulation

As described above, each node in a Boolean network model of transcriptional gene regulation usually corresponds to a gene, and each gene has incoming links from other genes that regulate its expression [6] (see Fig. 1(a) for a schematic illustration). Notably, in their network architecture, such Boolean modelings usually do not explicitly account for the transcription factors, the key molecules driving transcriptional regulation [37]. Consequently, Hannam *et al.* [37] proposed a bipartite Boolean network model that captures explicitly the two types of molecular species involved in transcriptional regulation namely, the genes (sequences of DNA) and TFs (see Fig. 1(b) for a schematic illustration).

The biological basis behind such a bipartite model of transcriptional regulation proposed by Hannam *et al.* [37] is as follows [37]. Firstly, a factor affecting a gene’s transcription rate can be either a single TF or a complex of TFs (e.g. heterodimer of two different TFs [49]). We refer to this factor as a transcriptional regulator (TR). Thus the presence of a transcriptional regulator may depend on the product of one or more genes. Secondly, multiple transcriptional regulators (TRs) can control the expression of a given gene. In the bipartite model, any species (i.e., genes or TRs, respectively) solely control the members of the other species (i.e., TRs or genes, respectively), and therefore, in the corresponging network representation there are no links between any two nodes of the same type.

Importantly, Hannam *et al.* [37] made a distinction between the case where a single TF acts as a TR of a target gene versus the case where a complex of two or more proteins acts as a TR of a target gene in the bipartite model. In the Results section, we expand the biological basis of this bipartite model to account for *cis*-regulatory modules such as enhancers as TRs involved in transcriptional regulation, and moreover, we evaluate the biological plausibility of composition structures using empirical data.

Formally, consider a subgraph in the bipartite network model of transcriptional regulation wherein a given gene has *r* incoming links from *r* TRs, that is, the expression of the given gene is directly controlled by *r* TRs. Furthermore, each of these *r* TRs in the subgraph have *t_i_* incoming links from *t_i_* genes where *i* ∈ [1,*r*], that is, each TR *i* is directly dependent on *t_i_* genes. In recent work, Fink and Hannam [41] termed such a subgraph in the bipartite model as a ‘composition structure’, and denoted it as {*t*_1_, *t*_2_,…,*t_r_*} (see Fig. 1(b)); since the composition graph is a tree of depth 2, the ordering of the degrees (i.e., *t_i_*s) is arbitrary and so one can force the sequence {*t*_1_, *t*_2_,…, *t_r_*} to be increasing. In their work, Fink and Hannam [41] assumed that the *t*_1_, *t*_2_,…,*t_r_* genes directly controlling the *r* TRs in the subgraph are distinct. Evidently, the sum *k* = *t*_1_ + *t*_2_ + … + *t_r_* gives the number of genes whose products directly regulate the targeted gene in the bipartite model. In other words, this sum *k* in the bipartite model gives the number of inputs *k* to a gene in the corresponding unipartite model.

Clearly, for a given value of *k*, there are multiple composition structures possible. For instance, the possible composition structures for *k* = 4 are: {1, 1, 1, 1}, {1, 1, 2}, {1, 3}, {2, 2} and {4}. Interestingly, Fink and Hannam [41] showed that in this bipartite network framework, certain composition structures can significantly restrict the possible set of logical update rules or BFs that can be assigned to genes. Fink and Hannam [41] refer to the subset of functions within all 2^2^*k*^^ BFs resulting from the restrictions imposed by the composition structure as “composed Boolean functions”. Next, we provide an overview of composed Boolean functions or “composed BFs”, and quantify the associated restriction.

### E. Composed Boolean functions

Consider a composition structure {*t*_1_, *t*_2_,…,*t_r_*} in the bipartite Boolean network framework. At the top thereof is a gene whose transcriptional regulation depends on the states of *r* TRs according to a BF *g* with *r* inputs. We denote the BF *g* as *g* = *g*(*y*_1_, *y*_2_,…, *y_r_*), where *y*_1_, *y*_2_,…,*y_r_* are the states of the *r* TRs. The state of each TR *i*, where *i* ∈ [1,*r*], in turn depends on the states of genes according to a BF *p_i_* with *t_i_* inputs.

Let us denote the states of the *k* = *t*_1_ + *t*_2_ + … + *t_r_* genes directly controlling the *r* TRs as *x*_1_,…, *x*_*t*_1__,…, *x*_*t*_1__+*t*_2_,…, *x_k_*. It follows that:

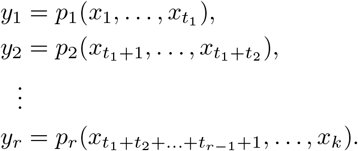

The regulation of a gene in the composition structure {*t*_1_, *t*_2_,…, *t_r_* } ultimately depends on the states of *k* genes according to some BF *h* of *k* inputs. This BF *h* is in fact the composition of the BFs *p*_1_, *p*_2_,…, *p_r_* fed into *g*, that is:

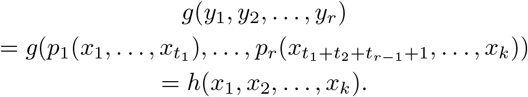

In the above equation, the BF *h* is said to be a composed BF. Note that there are no restrictions on the BFs that can be assigned to *p*_1_, *p*_2_,…, *p_r_* or *g*. Therefore, the upper limit on the possible number of composed BFs *h* is:

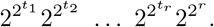

However, the 2^2*t*_1_^ 2^2*t*_2_^ … 2^2*t_r_*^ 2^2*^r^*^ BFs thereby composed are generally not all distinct, and it is necessary to remove the redundancies to obtain the set of (non-redundant) composed BFs. Such a non-redundant set of composed BFs is referred to as “biological logics” by Fink and Hannam [41], and in the present work we will refer to this non-redundant set of BFs as simply the “composed BFs”. From the definition of composed BFs, it follows that if a BF *h* is associated with a composition structure {*t*_1_, *t*_2_,…, *t_r_*}, then its complement 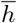 is also associated with the same composition structure (see Property A.1 in Appendix). Figure 1(b) provides a schematic illustration of a composed BF belonging to the composition structure {1, 2}.

For a given value of *k*, there are multiple composition structures {*t*_1_, *t*_2_,…, *t_r_*} such that *t*_1_ + *t*_2_ + … + *t_r_* = *k*. Fink and Hannam [41] have provided exact analytical expressions for the number of composed BFs in a composition structure. Following these analytical expressions, it can be easily shown that the composed BFs belonging to the two composition structures {1, 1, 1,…,1} and {*k*} do not restrict the space of *k*-input BFs (see Property A.2 in Appendix), they each include all 2^2*^k^*^ possible BFs. Thus, for *k*-input BFs, these two composition structures can be considered as trivial whereas the remaining composition structures are in fact non-trivial. Further, it is easy to see that there are no non-trivial composition structures for 1-input and 2-input BFs. Importantly, we excluded all the trivial composition structures from the analyses reported in this work, and in particular, we focus on non-trivial composition structures corresponding to 3, 4 and 5 input BFs.

## III. RESULTS

### A. Quantifying the presence of protein complexes that can act as transcriptional regulators

Hannam *et al.* [37] proposed to model transcriptional gene regulation via bipartite Boolean networks, thus allowing them to distinguish genes from proteins or protein complexes. Non-trivial composition structures then emerge if regulation requires complexes of TFs [37, 40] (see section II E). To our knowledge there has been no systematic study into such complexes in any organism, and we here attempt to fill this gap.

Genes often come in families following either segmental or whole genome duplications, and that is the case in particular for those coding for TFs. There are several organisms where it has been shown that the TFs within a given family form heterodimers or even multimers (and thus protein complexes) that contribute to gene regulation [50–53]. For instance the family of TFs called *auxin response factors* (ARFs) includes over 20 members in numerous plants and it has been shown that they form heterodimers that activate gene transcription [52, 54]. However, a quantitative assessment of the frequency at which heteromeric complexes contribute to gene regulation has not been carried out. The prevalence of such complexes in real-world gene regulatory networks can provide empirical support for the (frequent or not) occurrence of non-trivial composition structures. Here we estimate the number of heteromeric protein complexes contributing to gene regulation in *Homo sapiens*, while the case of *Saccharomyces cerevisiae* is treated in Appendix B.

First, we obtained a list of 1325 macromolecular complexes in *H. sapiens* from the EBI Complex Portal database [55], and the list of 1639 human TFs provided by Lambert *et al.* [56] (see http://humantfs.ccbr.utoronto.ca/). Among the 1639 human TFs, we selected only those TFs that were reviewed in the SWISS-PROT protein database, resulting in a list of 1617 human TFs that was used for further analysis. We found that among the 1325 complexes in *H. sapiens*, 169 satisfy the constraint of being heteromeric with all subunits corresponding to TFs (Supplementary Table S1). Of those, 165 are heterodimers and the remaining 4 are heterotrimers. Furthermore, there are 84 unique TFs composing these 169 complexes. Second, we manually searched for DNA binding evidence for each of these 169 heteromeric complexes and found that DNA binding has been verified for 86 of them. This then leaves us with 86 validated complexes of TFs that act as TRs, and thus, are likely candidates for forming composition structures.

We also explored the specific importance of heteromers of TFs belonging to particular families. Indeed, it is known that certain classes of TFs, for instance basic leucine zipper (bZIP) [57, 58] and basic helix-loop-helix (bHLH) [59, 60] classes, tend to bind to DNA as homo- or hetero-dimers. Knowing the prevalence of such TFs could shed light on the abundance of dimeric complexes which act as TRs. We thus determined the TF families within 1617 complexes in *H. sapiens* using the JASPAR database [61] which provides a manually curated list of DNA transcription factor binding motifs, the corresponding TFs, as well as their classes. Among the 1617 TFs in H. *sapiens,* we counted the ones that belong to the bZIP and bHLH classes and found 36 TFs that belong to the bZIP class and 38 TFs that belong to the bHLH class (see Supplementary Table S2).

### B. Composition structures arising through enhancers

Bipartite Boolean network models provide a quite general framework and so for instance composition structures can accommodate other mechanisms of eukaryotic gene regulation than the one involving complexes as covered in the previous sub-section. Here, we propose one such alternative picture where the intermediate transcriptional regulators (TRs) are no longer protein complexes but are associated with *cis*-regulatory modules such as promoters, enhancers, or insulators. In eukaryotes, transcription is typically regulated via the binding of TFs upstream of the gene [43, 44, 62]. Promoters are located close to the transcription start site where RNA polymerases and transcription factors assemble to initiate transcription [63]. Enhancers on the other hand may be located at rather large distances (in fact both upstream or downstream) of the target gene they regulate [64]. Enhancers are “active” or “inactive” based on whether their chromatin state is accessible or not; in the former case, transcription factor binding sites within these enhancers can attract specific TFs and thus modulate transcription of nearby genes [44]. Interestingly, a given enhancer typically contains multiple such binding sites and is thus considered to be a *cis*-regulatory module [44, 65].

Fig. 2(a) and Fig. 2(b) illustrate how enhancers and promoters may act as TRs in the composition structure {2, 3} where we have chosen to have 2 TFs binding to the promoter and 3 TFs binding to the enhancer. One can suppose that an abundance of enhancers containing multiple TF binding sites is suggestive of the prevalence of non-trivial composition structures in real-world gene regulatory networks. In view of this possibility, we perform an analysis to provide a quantitative estimate of the number of TFs that bind to active enhancers in two widely-studied human cell lines namely, HepG2 and K562.

**FIG. 2.**
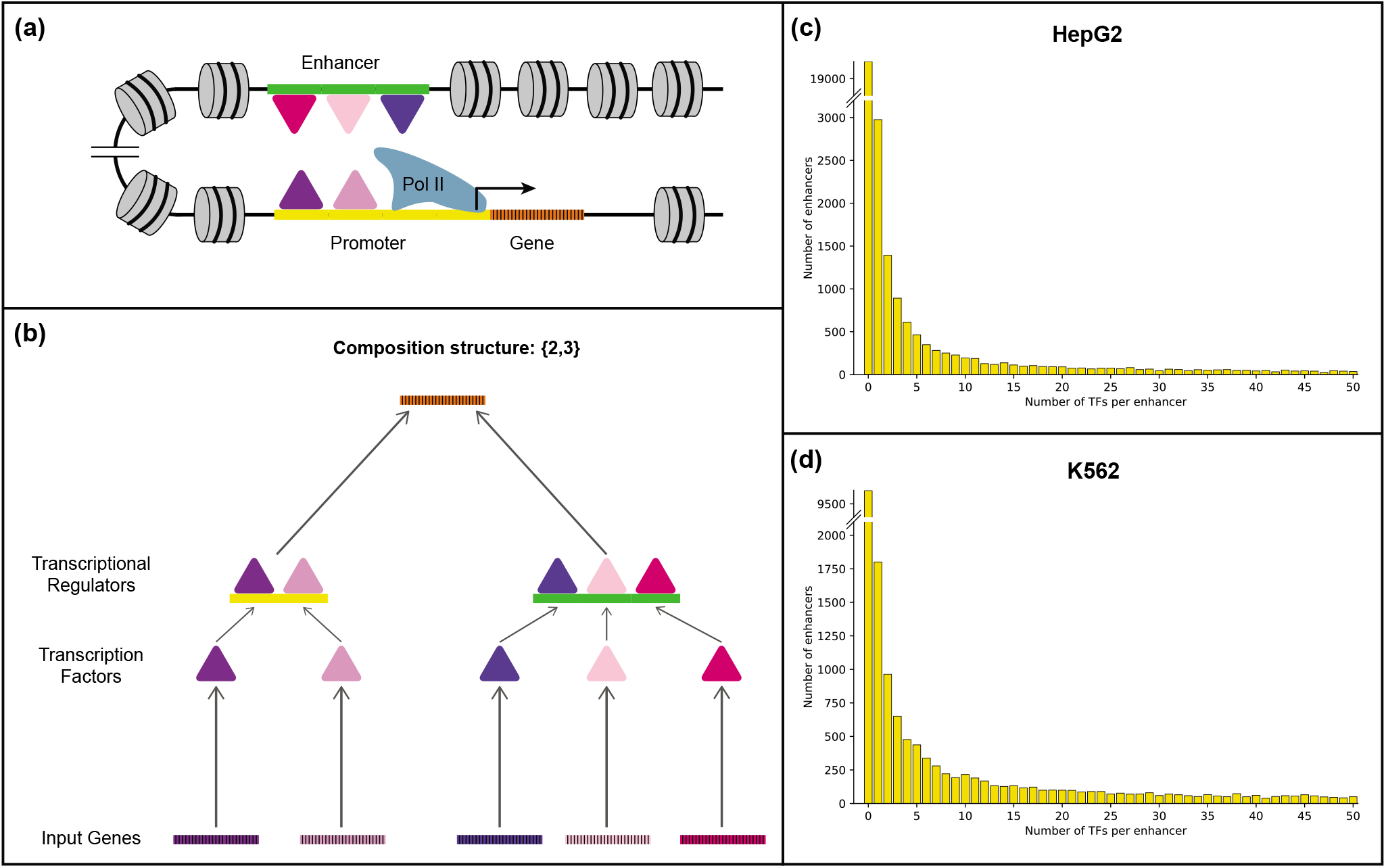
Non-trivial composition structures arising due to enhancers bound by multiple transcription factors. (a) A biologically plausible mechanism revealing the occurrence of non-trivial composition structures in transcriptional gene regulation. Multiple transcription factors (TFs) can bind to the promoter as well as the enhancer region(s) of a target gene. The enhancers and promoters bound by TFs then act as transcriptional regulators (TRs) of their target genes, resulting in non-trivial composition structures. (b) A schematic representation of the composition structure {2, 3} arising in subfigure (a). The target gene is regulated by an active promoter that is bound by 2 TFs, and an active enhancer that is bound by 3 TFs. (c) Bar plot showing the number of active enhancers bound by a given number of TFs in the HepG2 cell line in humans. We found that 32.68% of the active enhancers in HepG2 are bound by at least 2 TFs. (d) Bar plot showing the number of active enhancers bound by a given number of TFs in the K562 cell line in humans. We found that 44.31% of the active enhancers in K562 are bound by at least 2 TFs. These results suggest that non-trivial composition structures are prevalent in gene regulatory networks.

To perform our analyses, we relied on two published datasets: (i) TF binding regions, identified from ChIP-seq experiments, and (ii) active enhancers. We focused on the two well-studied human cell lines HepG2 and K562 because they are rich in such data. We extracted the TF binding regions from the ChIP-seq data in the human ENCODE project [66] and the active enhancers from data processed using the STARRPeaker peak-calling software [67]. In particular, the ChIP-seq narrowPeak bed files for HepG2 and K562 cell lines were last downloaded on 28^th^ April 2022 and 29^th^ April 2022, respectively from the human ENCODE project website: https://www.encodeproject.org. We then considered that a TF binds to an active enhancer if and only if both the midpoint and the summit of the ChIP-seq peaks for that TF fall within the active enhancer region. The processed datasets from ENCODE used for this analysis and the associated codes are available at: https://github.com/asamallab/CoSt.

For the cell line HepG2, we used ChIP-seq peaks for 458 unique TFs and there are a total of 32929 active enhancers. 2976 enhancers had exactly one TF binding within their region while 10754 enhancers had two or more TFs binding within their region, representing 32.68% of the total number of enhancers in HepG2 (Fig. 2(c)). Additionally, of the 458 TFs for which data is available in HepG2, we found that 456 TFs bind to at least one of the enhancers detected as active. For the cell line K562, we used ChIP-seq peaks for 323 unique TFs and there are a total of 20471 active enhancers. 1801 enhancers had exactly one TF binding within their region while 9071 enhancers had two or more TFs binding within their region, representing 44.31% of the total number of enhancers in K562 (Fig. 2(d)). Additionally, of the 323 TFs for which data is available in K562, we found that 322 TFs bind to at least one of the enhancers detected as active. The fact that 32.68% and 44.31% of the active enhancers in HepG2 and K562, respectively can be bound by at least two TFs suggest that non-trivial composition structures indeed do arise frequently in gene regulatory logics.

### C. Accounting for all possible permutations of the inputs in a composition structure

In their procedure to count BFs arising from a composition structure, Fink and Hannam [41] do not account for permutations of the input variables, that is they ignore the labels of the inputs. To illustrate this, consider a composed BF of the type *g*(*p*_1_(*x*_1_),*p*_2_(*x*_2_, *x*_3_)) that belongs to the composition structure {1, 2} and corresponds to a 3-input BF *h*(*x*_1_, *x*_2_, *x*_3_). Taking *p*_1_(*x*_1_) = *x*_1_, *p*_2_(*x*_2_,*x*_3_) = *x*_2_ *x*_3_, and *g*(*x,y*) = *x* + *y* leads to the composed BF *h*(*x*_1_, *x*_2_, *x*_3_) = *x*_1_ + *x*_2_ *x*_3_. However the BFs obtained by permuting the labels of these variables, namely *x*_2_ + *x*_1_ *x*_3_ and *x*_3_ + *x*_1_ *x*_2_, are just as relevant biologically; indeed, the labels point to genes and these are hardly ever equivalent. Thus, we count all three of the cases above as valid composed BFs. In contrast, Fink and Hannam [41] count them as one composed BF. Note that this example shows that the two ways of counting are not generally related by the number of permutations (*k*!) of *k* labels because of possible symmetries within these expressions.

Table 1 provides a comparison of the number of distinct BFs allowed by different composition structures for *k* ≤ 5 inputs both with and without including all possible permutations of the input variables. Table 1 also provides these results as fractions among all possible BFs for *k* ≤ 5 inputs. Naturally, we find that accounting for all possible permutations of inputs increases the number of BFs in a composition structure in comparison to those reported by Fink and Hannam [41]. However, this does not alter the central result of Fink and Hannam [41], that is, composition structures significantly restrict the space of possible BFs. This is evident from the trends for the fractions of composed BFs among all possible BFs as a function of the number of inputs (see Table 1). Moreover, including all possible permutations of inputs ensures that all isomorphisms of a BF in a composition structure are present therein (see section II B).

**TABLE 1.** Comparison of the number and fraction of BFs allowed by different composition structures, with and without including all possible permutations of input variables. The composition structures in a bipartite Boolean network model of transcriptional gene regulation are categorized based on the number of inputs *k* to a gene in the corresponding unipartite Boolean network model. The column “Number of composed BFs” gives the number of distinct BFs in a composition structure, and the subcolumns provide a comparison of the number of such BFs both without and with the accounting for all possible permutations of the input variables. The column “Fraction of composed BFs” gives the fraction of distinct BFs in a composition structure among all possible BFs for a given number *k* of inputs, and the subcolumns provide a comparison of the fraction of such BFs both without and with the accounting for all possible permutations of the input variables.

| Inputs<br>($k$) | Composition<br>structure | Number of composed BF's | | Fraction of composed BF's | |
| --- | --- | --- | --- | --- | --- |
|  |  | without<br>permutation | with<br>permutation | without<br>permutation | with<br>permutation |
| 1 | {1} | 4 | 4 | 1 | 1 |
| 2 | {2} | 16 | 16 | 1 | 1 |
|  | {1,1} | 16 | 16 | 1 | 1 |
| 3 | {3} | 256 | 256 | 1 | 1 |
|  | {1,2} | 88 | 152 | 0.344 | 0.594 |
|  | {1,1,1} | 256 | 256 | 1 | 1 |
| 4 | {4} | 65536 | 65536 | 1 | 1 |
|  | {1,3} | 1528 | 4864 | 0.023 | 0.074 |
|  | {2,2} | 520 | 1208 | 0.008 | 0.018 |
|  | {1,1,2} | 1696 | 6216 | 0.026 | 0.095 |
|  | {1,1,1,1} | 65536 | 65536 | 1 | 1 |
| 5 | {5} | 4294967296 | 4294967296 | 1 | 1 |
| | {1,4} | 393208 | 1921928 | $9.16 \times 10^{-05}$ | $4.47 \times 10^{-04}$ |
| | {2,3} | 9160 | 71608 | $2.13 \times 10^{-06}$ | $1.67 \times 10^{-05}$ |
| | {1,1,3} | 30496 | 263488 | $7.10 \times 10^{-06}$ | $6.13 \times 10^{-05}$ |
| | {1,2,2} | 11344 | 100768 | $2.64 \times 10^{-06}$ | $2.35 \times 10^{-05}$ |
| | {1,1,1,2} | 457216 | 3446488 | $1.06 \times 10^{-04}$ | $8.02 \times 10^{-04}$ |
|  | {1,1,1,1,1} | 4294967296 | 4294967296 | 1 | 1 |

### D. Overlap of Boolean functions across various *k*-input composition structures

There are multiple composition structures {*t*_1_,…, *t_r_*} possible for a given number of inputs *k* such that *t*_1_ + *t*_2_ +… + *t_r_* = *k*, and each composition structure allows a certain set of BFs. However, composed BFs can belong to more than one composition structure. Therefore, it is worthwhile to examine the overlaps of composed BFs across all non-trivial composition structures with a given number of inputs *k*. Here, we analysed the intersections of composed BFs across non-trivial composition structures for *k* = 4 and *k* = 5 inputs. We reiterate that there are no non-trivial composition structures for *k* = 1 and *k* = 2 inputs, and note that {1,2} is the only non-trivial composition structure for *k* = 3.

For *k* = 4 inputs, we find that the set of BFs in the composition structure {2, 2} is a proper subset of the set of BFs in {1, 1, 2} (see Figure 3(a)). For *k* = 5 inputs, we find that the set of BFs in the composition structure {2, 3} is a proper subset of the set of BFs in {1, 1, 3} as well as {1, 1, 1, 2}, and the set of BFs in the composition structure {1, 2, 2} is a proper subset of the set of BFs in {1, 1, 1, 2} (see Figure 3(a)). Further, we give the number of BFs in all possible intersections of non-trivial composition structures for *k* = 4 and *k* = 5 inputs through UpSet plots [68] in Figure 3(b) and (c), respectively.

**FIG. 3.**
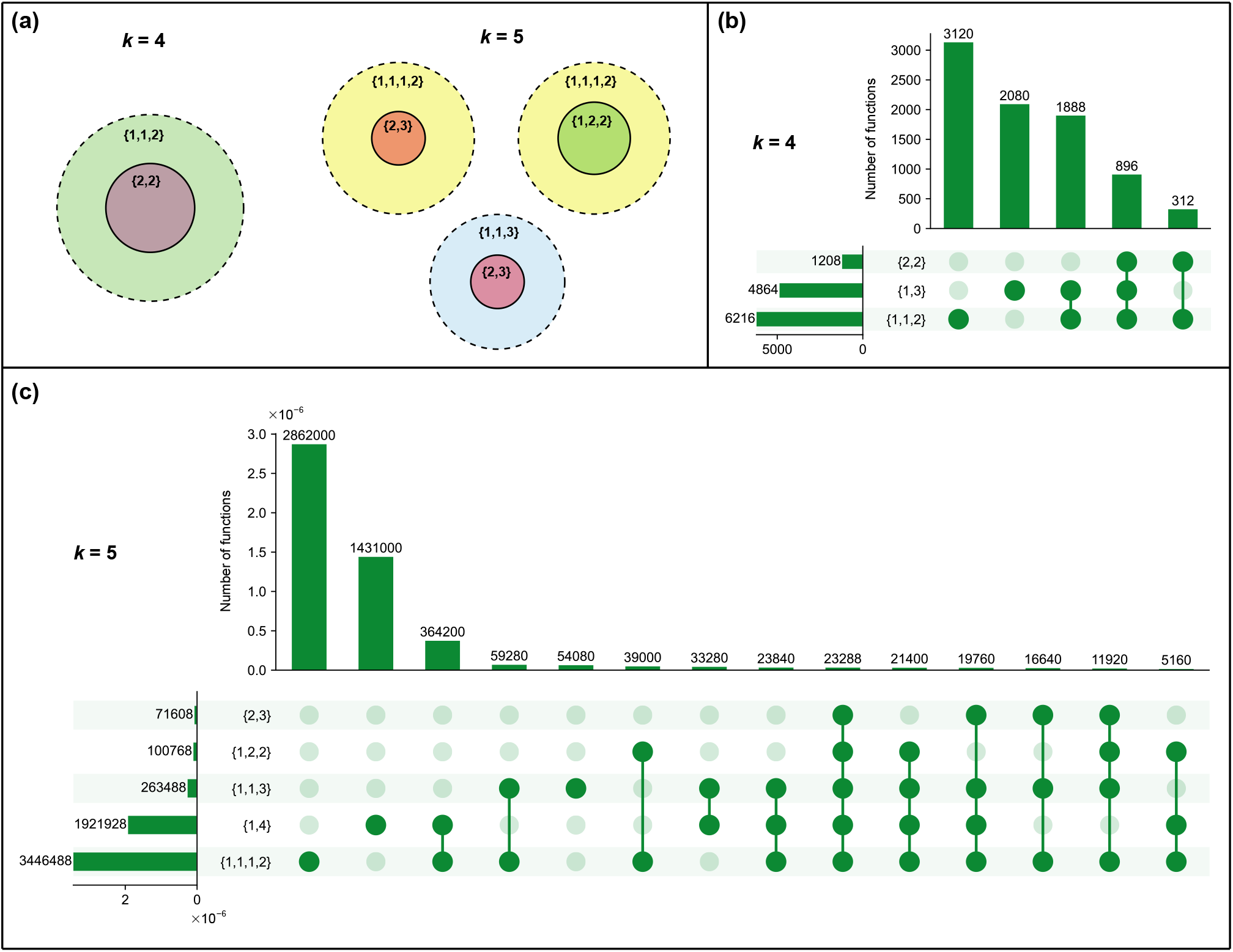
Overlaps between the sets of BFs compatible with different composition structures at *k* = 4 and *k* = 5 inputs. (a) Venn diagrams illustrating *proper* subsets among the sets of non-trivial composition structures at *k* = 4 and *k* = 5 inputs. (b) UpSet plot illustrating the number of BFs that are present in all possible intersections of non-trivial composition structures at *k* = 4 inputs. (c) UpSet plot illustrating the number of BFs that are present in all possible intersections of non-trivial composition structures at *k* = 5 inputs. The horizontal bars in the UpSet plots indicate the number of BFs that are present in different composition structures. The vertical bars indicate the number of BFs that are simultaneously present in some and absent from other composition structures, as specified by the underlying dark and light green circles.

### E. Comparing restriction levels: composition structures vs biologically meaningful types

Clearly, imposing a non-trivial composition structure significantly restricts the space of allowed BFs within the complete space of BFs with *k* inputs. As shown by some of us recently [23], the same holds when imposing certain biologically meaningful properties. Here, we compare the level of restriction achieved by four established biologically meaningful types of BFs, namely UFs, CFs, NCFs and RoFs, to that achieved by composed BFs of a given composition structure, in the space of all BFs with *k* inputs. Among the four different types of biologically meaningful BFs, it is known that the NCFs represent the smallest fraction in the space of all BFs [23]. For *k* = 4 and *k* = 5 inputs, we find that certain composition structures restrict very strongly, though less than NCFs (see Supplementary Table S5). Specifically, at *k* = 4 inputs, {2, 2} is the most restrictive one. The composed BFs in {2, 2} occupy a fraction of 0.018 among all BFs, which is 1.63 times greater than the fraction occupied by NCFs at *k* = 4 (whose value is 0.011). For *k* = 5 inputs, {2, 3} is the most restrictive composition structure. The BFs in {2, 3} occupy a fraction of 1.67 × 10^-5^, which is about 6.76 times greater than the fraction occupied by NCFs at *k* = 5 (whose value is 2.47 × 10^-6^). In Supplementary Table S5, we compare the fraction of BFs in the most restrictive composition structure to the fractions for each of the four types of biologically meaningful BFs for *k* ≤ 5 inputs.

We next evaluated how often a BF in a composition structure also displays biologically meaningful properties. Table 2 shows the number of composed BFs that belong to each of the four types of biologically meaningful BFs for non-trivial composition structures with *k* ≤ 5 inputs. Clearly imposing BFs to be biologically meaningful and to be compatible with a given composition structure severely restricts the possible BFs. We also find that certain types of biologically meaningful BFs, in particular NCFs, are proper subsets of BFs in certain composition structures. Specifically, all the 64 NCFs with *k* = 3 inputs are contained in the composition structure {1, 2}, all the 736 NCFs with *k* = 4 inputs are contained in the composition structures {1, 3} and {1, 1, 2}, and all the 10624 NCFs with *k* = 5 inputs are contained in the composition structures {1, 4}, {1, 1, 3} and {1, 1, 1, 2}. Moreover, all CFs with *k* = 3, 4, and 5 inputs are a subset of the composition structures {1, 2}, {1, 3} and {1, 4}, respectively, whereas all RoFs with *k* = 4 and 5 inputs are a subset of the composition structures {1, 1, 2} and {1, 1, 1, 2}, respectively. In Supplementary Table S6, we provide the fraction of composed BFs that belong to each of the four types of biologically meaningful BFs for non-trivial composition structures with *k* ≤ 5 inputs.

**TABLE 2.** Number of BFs in different composition structures that display biologically meaningful properties. The number of BFs within a non-trivial composition structure that also belong to each of the four types of biologically meaningful functions, namely Unate functions (UF), Canalyzing functions (CF), Nested canalyzing functions (NCF) and Read-once functions (RoFs). The column ‘Number of composed BFs’ gives the number of BFs that are allowed in a given composition structure.

| Composition structure | Number of composed BFs | Number of biologically meaningful BFs in composition structure |  |  |  |
| --- | --- | --- | --- | --- | --- |
|  |  | UF | CF | NCF | RoF |
| {1,2} | 152 | 96 | 120 | 64 | 64 |
| {1,3} | 4864 | 1210 | 3514 | 736 | 736 |
| {2,2} | 1208 | 634 | 730 | 224 | 320 |
| {1,1,2} | 6216 | 1370 | 1850 | 736 | 832 |
| {1,4} | 1921928 | 41676 | 1292276 | 10624 | 12544 |
| {2,3} | 71608 | 13676 | 33596 | 3264 | 6784 |
| {1,1,3} | 263488 | 26156 | 80996 | 10624 | 14144 |
| {1,2,2} | 100768 | 17836 | 25236 | 5504 | 9984 |
| {1,1,1,2} | 3446488 | 61516 | 122516 | 10624 | 15104 |

We also computed the number and fraction of composed BFs for different composition structures which have odd bias. Recently, some of us showed that BFs with odd bias are preponderant among BFs in reconstructed Boolean network models of biological systems [23]. Furthermore, it was shown that NCFs [69] and RoFs [23] have odd bias. Here, we find that the fraction of BFs with odd bias in any composition structure with *k* ≤ 5 inputs is less than 0.5 (see Supplementary Table S7). Additionally, we find that BFs – with any given even bias – occur in all composition structures with *k* ≤ 5 inputs. In Supplementary Table S7, we list the odd biases of BFs that are present in composition structures with *k* ≤ 5 inputs.

### F. Enrichments of composed BFs in reconstructed biological networks

In this section, we present the results of our analyses of the abundances of composed BFs in a compiled reference biological dataset of 2687 BFs from 88 published Boolean network models of biological systems [23]. In Appendix C, we provide details about the compilation and curation of this empirical dataset.

To begin, we computed two proportions for each possible composition structure. The first is the proportion of BFs with *k* inputs in the reconstructed biological networks that belong to the given composition structure (bar plots in Figure 4(a)). The second is the corresponding proportion in the “random ensemble” with *k* inputs; that proportion is thus given by the number of BFs with *k* inputs that are compatible with the given composition structure, divided by the total number of BFs (black dots in Figure 4(a)). The results show that the proportions of composed BFs in the *reference biological dataset* are larger than in the ensemble of random BFs, for each composition structure, indicating that non-trivial composed BFs are enriched in real biological networks.

**FIG. 4.**
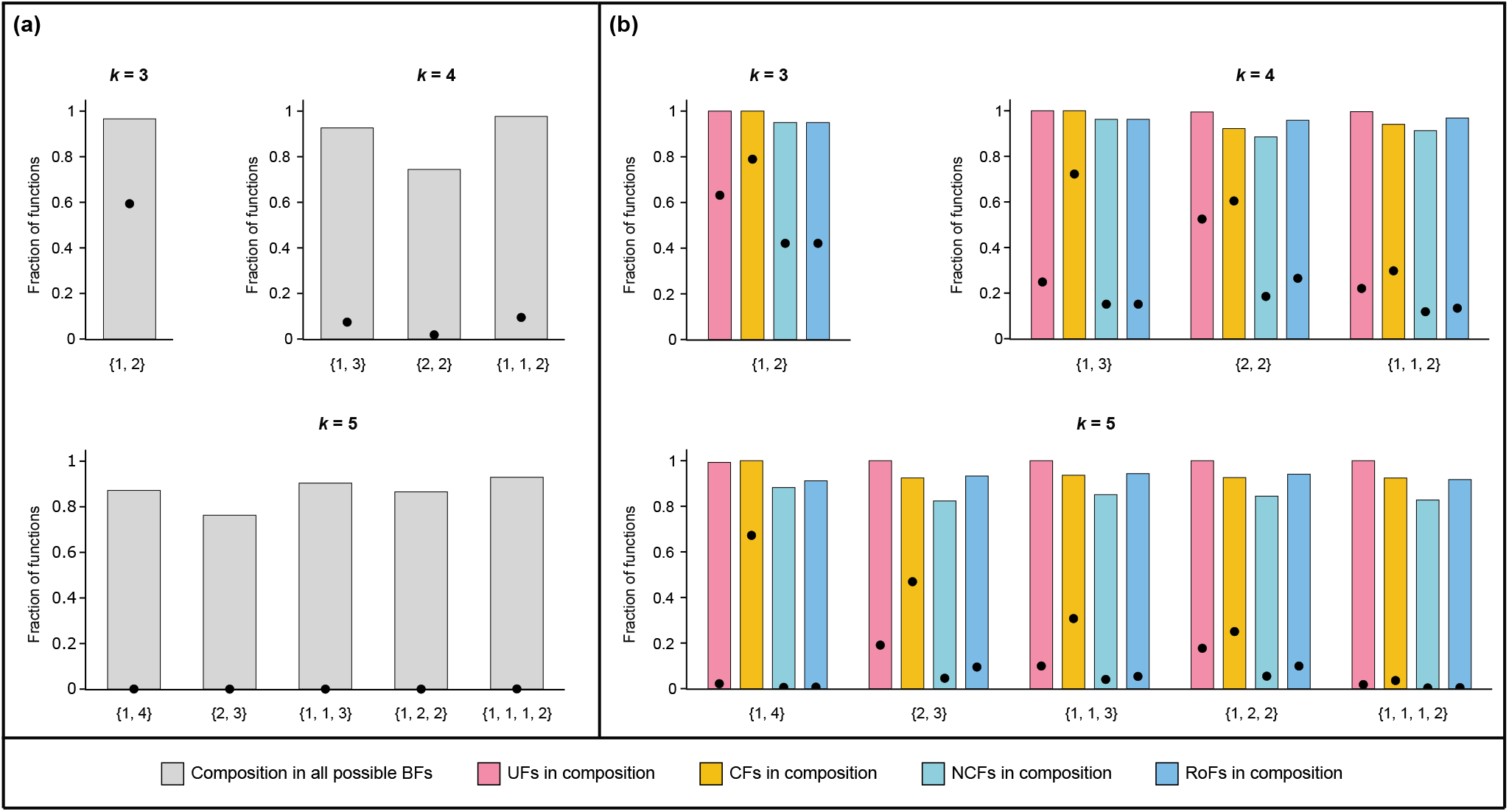
Abundance of composed BFs in reconstructed biological networks. (a) Bar plots giving the fraction of BFs in the reference biological dataset that are compatible with each of the composition structures. The black dots indicate the fraction when considering all possible BFs instead of only the ones in the reference biological dataset. (b) For all BFs of the reference biological dataset compatible with a given composition structure, the bars give the fraction of these BFs that belong to each of the four biologically meaningful sub-types: Unate functions (UF), Canalyzing functions (CF), Nested canalyzing functions (NCF), and Read-once functions (RoFs). Again, the black dots give these fractions when considering instead all possible BFs.

To consider this question in greater depth, we define the “enrichment factor” as the ratio of the first and the second proportions. From Figure 4(a) we can see that the enrichment factors are very large for *k* = 4 and 5. For instance, for the composition structures {2, 2} and {2, 3} that are the most restrictive composition structures for *k* = 4 and *k* = 5 inputs, the corresponding enrichment factors are 40.37 and 45760.08. To check the level of significance of this effect, we applied a standard statistical test (see Appendix D). In Supplementary Table S8, we list the enrichment factors for all non-trivial composition structures having *k* ≤ 5 inputs and we give the corresponding one-sided *p*-values. These *p*-values show that the enrichment effects are indeed statistically significant, providing evidence in biological systems of a selection pressure in favor of each of the non-trivial composition structures.

To refine the analysis, we ask whether functions in the reference biological dataset compatible with a given composition structure might be further enriched for a biologically meaningful sub-type of BFs. To find out, we introduce the *relative enrichment E_R_* for a given sub-type *s* of biologically meaningful BFs and a given composition structure. *E_R_* is defined as a ratio where the numerator is the proportion of sub-type *s* BFs observed among the BFs arising in the reference biological dataset that are also compatible with the given composition structure, and the denominator is the analogous proportion when considering all BFs rather than the ones in the biological dataset. The denominator is thus the proportion of sub-type *s* BFs observed among all BFs compatible with the given composition structure. See Appendix D for the detailed methodology. If *s* plays no biological role, we expect *E_R_* to be around 1. In contrast, large values for *E_R_* are indicative of a selection pressure for *s* in addition to possible selection pressures on composition structures.

Figure 4(b) is a bar plot of the fractions in the reference biological dataset of the four biologically meaningful subtypes when focusing on the BFs satisfying a given composition structure. In addition, the black dots give the corresponding fractions when using the random ensemble instead of the reference biological dataset. Clearly the different enrichments *E_R_* are larger than 1 for all non-trivial composition structures with number of inputs *k* ≤ 5, and this suggests that the four biologically meaningful subtypes of composed BFs are enriched among composition structures in the reference biological dataset. Table 3 gives the *E_R_* values for the four biologically meaningful sub-types in non-trivial composition structures with number of inputs *k* ≤ 5. Furthermore, the computed relative enrichment values are statistically significant as determined by one-sided *p*-values (see Supplementary Table S9).

**TABLE 3.**
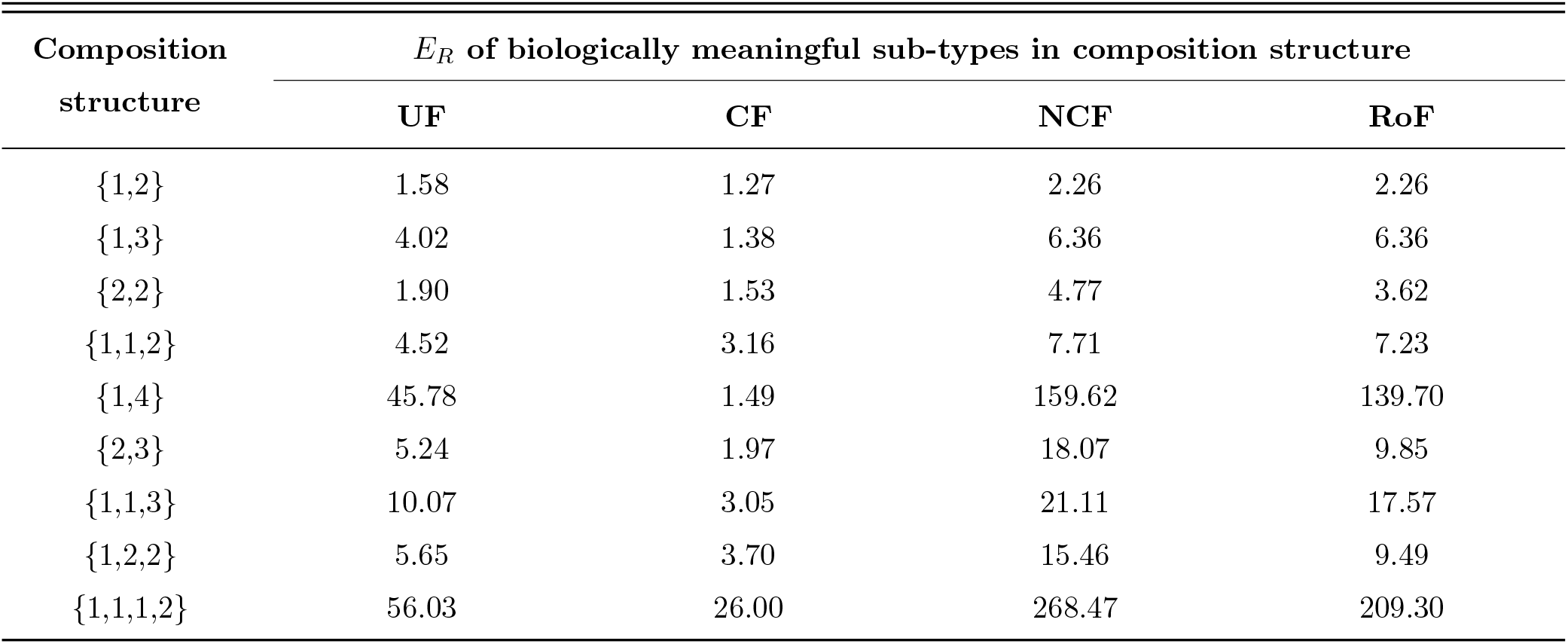
Relative enrichment of biologically meaningful BFs among composed BFs of different composition structures in the reference biological dataset. This table gives the relative enrichment values *E_R_* in the reference biological dataset for the four biologically meaningful sub-types within composed BFs for different non-trivial composition structures with number of inputs *k* ≤ 5. These four biologically meaningful sub-types within composed BFs include those BFs in a composition structure that also happen to be Unate functions (UF), Canalyzing functions (CF), Nested canalyzing functions (NCF), or Read-once functions (RoFs).

A previous analysis [23] showed that biologically meaningful BFs are enriched in our reference biological dataset. Notably, those enrichments are likely driven by complexity minimization, with NCFs and RoFs respectively minimizing two complexity measures namely, average sensitivity and Boolean complexity [23]. An immediate question that then arises is whether the enrichments of composed BFs as found in Fig. 4 might just be driven by enrichments of NCFs and RoFs. To examine that possibility, let *T_C_* denote the set of BFs allowed by a composition structure *C* at a given number of inputs *k*, and let *T_NCF_* denote the set of NCFs with *k* inputs. We have determined the enrichment factors of three disjoint sets of BFs: composed BFs that are also NCFs (i.e., *T_C_* ⋂*T_NCF_*), composed BFs that are not NCFs (i.e., *T_C_* \ *T_NCF_*), and NCFs that are not composed BFs (i.e., *T_NCF_* \ *T_C_*). Table 4 shows the enrichment factors for these three disjoint sets of BFs, for all non-trivial composition structures with *k* ≤ 5 inputs. We find that the BFs belonging to the set *T_C_*⋂*T_NCF_* display a very high enrichment factor. Moreover, for composition structures {2, 2}, {2, 3} and {1, 2, 2}, we find that both the sets *T_C_* \*T_NCF_* and *T_NCF_* \ *T_C_* are enriched in the biological datasets. However, the enrichment factor is much larger for the set *T_NCF_*\*T_C_*. Finally, for the composition structures {1, 2}, {1, 3}, {1, 1, 2}, {1, 4}, {1, 1, 3} and {1, 1, 1, 2} that are a superset of the corresponding NCFs, we find that the set *T_C_* \ *T_NCF_* is either depleted or shows a lower enrichment factor compared to the set *T_C_* ⋂ *T_NCF_*. After repeating the above analysis for RoFs to estimate the enrichment factors for *T_C_* ⋂ *T_RoF_*, *T_C_* \ *T_RoF_* and *T_RoF_* \ *T_C_*, we find that the results are similar to that for NCFs (Table 4). Furthermore, all these enrichment factors are statistically significant as determined by one-sided *p*-values (Supplementary Table S10). These results suggest that although composed BFs are subject to positive selection in real biological networks, the primary driving force for enrichment is the property of being an NCF or an RoF.

**TABLE 4.** Comparison between the enrichments of composed BFs and biologically meaningful BFs of minimum complexity in the reference biological dataset. The table provides the enrichment factors when composed BFs in nontrivial composition structures with *k* ≤ 5 inputs are compared with two biologically meaningful BFs of minimum complexity namely, nested canalyzing functions (NCFs) and Read-once functions (RoFs). *T_C_* denotes the set of composed BFs allowed by a composition structure at a given number of inputs *k, T_NCF_* denotes the set of all *k*-input NCFs, and *T_RoF_* denotes the set of all *k*-input RoFs. ⋂ represents the intersection of two sets and \ represents the set-theoretic difference. “–” in the columns *T_NCF_* \ *T_C_* or *T_RoF_* \ *T_C_* indicates that the NCFs or RoFs are a subset of the set of BFs allowed by the composition structure.

| Composition<br>structure | $T_C \cap T_{NCF}$ | $T_C \setminus T_{NCF}$ | $T_{NCF} \setminus T_C$ | $T_C \cap T_{RoF}$ | $T_C \setminus T_{RoF}$ | $T_{RoF} \setminus T_C$ |
| --- | --- | --- | --- | --- | --- | --- |
| {1,2} | 3.67 | 0.14 | – | 3.67 | 0.14 | – |
| {1,3} | 79.38 | 0.55 | – | 79.38 | 0.55 | 37.04 |
| {2,2} | 192.78 | 5.68 | 29.77 | 146.06 | 2.29 | 29.77 |
| {1,1,2} | 79.38 | 1.02 | – | 74.49 | 0.38 | – |
| {1,4} | 310977.13 | 230.48 | – | 272157.87 | 173.03 | 96791.63 |
| {2,3} | 826630.05 | 8459.68 | 82296.27 | 450476.77 | 3397.73 | 72800.54 |
| {1,1,3} | 310977.13 | 2286.48 | – | 258889.63 | 883.34 | 0.00 |
| {1,2,2} | 570245.27 | 6069.12 | 32263.88 | 350214.73 | 2426.14 | 32263.88 |
| {1,1,1,2} | 310977.13 | 200.33 | – | 242434.78 | 96.28 | – |

We have also examined these questions for the other sub-types of biologically meaningful BFs. Supplementary Table S11 lists the corresponding enrichment factors while Supplementary Table S12 lists the associated *p*-values. First, we find that the set of BFs that are UFs but not composed BFs (i.e., *T_UF_* \ *T_C_*) are enriched whereas those BFs that are composed BFs but not UFs (i.e., *T_C_* \ *T_UF_*) are highly depleted. This suggests that UFs could also be a possible driving factor for the enrichment of composed BFs in biological networks. Second, the set of BFs that are composed BFs but not CFs (i.e., *T_C_* \ *T_CF_*) are highly enriched compared to the set of BFs that are CFs but not composed BFs (i.e., *T_CF_* \ *T_C_*). These results provide evidence for composition structures as a driving factor for the enrichment of CFs in real biological networks where NCFs and RoFs themselves are not significant drivers of such enrichments.

## IV. DISCUSSION AND CONCLUSIONS

We began our empirical study into the potential biological relevance of non-trivial composition structures arising in bipartite gene regulatory networks by investigating two different scenarios. In the first we estimated the degree of occurrence of heteromeric complexes formed by DNA-binding proteins while in the second we characterized co-occurrences of TF binding sites in enhancers. Recall that a composition is called non-trivial when the number of terms in the composition structure is greater than 1 and at least one of its entries is not equal to 1 (see section II E).

In the scenario of transcriptional regulation by heteromeric complexes as proposed by Hannam *et al.* [37], a non-trivial composition structure arises when a gene is regulated by at least two transcriptional regulators, of which at least one is a heteromeric protein complex made up of at least two monomers. Composition structures can therefore be important if a substantial fraction of genes are transcriptionally regulated by protein complexes. Generally that will require many different such complexes, but one cannot exclude that just a few complexes are involved in the control of many genes. From the data on macromolecular complexes in humans obtained from the EBI Complex Portal [55], we find that for approximately 6.5% of the complexes (86 out of 1325), all of their monomeric subunits are identified as TFs. (For such an identification, we imposed that they be present in the database of 1617 human TFs from Lambert *et al.* [56] and come with strong evidence for DNA binding as ascertained by manual curation of the literature.) Furthermore, we find that 4.57% of the human TFs belong to the bZIP and bHLH classes that are known to bind to DNA as homodimers or heterodimers.

It is likely that the collection of complexes in the EBI Complex Portal are biased towards complexes which do not act as TRs given that the detection and characterization of heteromeric protein complexes which act as TRs is experimentally challenging. Though our empirical analysis provides some support for Hannam’s picture of heteromeric protein complexes acting as TRs, the existing data on such complexes is insufficient to quantitatively estimate the prevalence of composition structures in real-world gene regulatory networks. Another point that requires critical assessment in this picture of gene regulation is the number of logic rules that govern the formation of the heteromeric complexes. Since a heteromeric complex is a conjunction of all its monomeric subunits, the only Boolean logic rule which captures the formation of a complex is the one linking all the components by the “AND” operator. In a general bipartite Boolean network, the upper limit for the number of logics possible for the composition structure {*t*_1_, *t*_2_,…, *t_r_*} is 2^2*t*_1_^ 2^2*t*_2_^ … 2^2*t_r_*^ 2^2*^r^*^, whereas if one imposes the “AND” logic for the formation of protein complexes only 2^2*^r^*^ logics are possible.

The flexibility of the bipartite formalism allows us to capture a more nuanced scenario in gene regulation that involves *cis*-regulatory elements (such as enhancers and promoters) and the transcription factors (TFs) which bind to them. In our picture, a target gene is regulated by *cis*-regulatory elements which act as transcriptional regulators (TRs) and each *cis*-regulatory element acting in a way that depends on the TFs that bind therein (see Fig. 2(a)). Thus, a non-trivial composition structure is realized when a gene is regulated by at least two *cis*-regulatory elements, one of which is regulated by at least two TFs. We thus inferred whether non-trivial composition structures of this kind arise in gene regulatory networks by determining how often the enhancers of a gene are bound by at least two TFs. By analyzing ChIP-seq and enhancer datasets in the two human cell lines HepG2 and K562, we find that 32.68% and 44.31% of their respective active enhancers bind to at least two TFs. Our result suggests that composition structures with *cis*-regulatory elements acting as transcriptional regulators are likely to be prevalent in bipartite gene regulatory networks. We remark that experimental limitations do not allow for the detection of all the enhancers for a given target gene in a given cell type, preventing the identification of exact composition structures from empirical data.

Fink and Hannam [41] showed that composition structures can severely restrict the number of Boolean logics in the space of all BFs. In this contribution, we address many questions from the perspective of providing a comprehensive comparison between the BFs of a composition structure and the biologically meaningful BFs. The main questions we address are as follows: (i) How restrictive are composition structures compared to biologically meaningful logic rules? (ii) For a given number of inputs, do the BFs belonging to two different composition structures overlap? (iii) Do BFs in a given composition structure overlap with that of the biologically meaningful logics? (iv) Are composed BFs enriched in an empirical dataset of biological logic rules? First, we provide the corrected values for the number of BFs belonging to a given composition structure by accounting for all the isomorphisms for each of the composed BFs (Fink and Hannam [41] leave these out of their work). We then compare the degree of restrictiveness imposed by composition structures vis-a-vis biologically meaningful logic rules and find that for all inputs up to 5-input BFs, the NCFs and RoFs are more restrictive than the most restrictive composition structures. Next, we quantify the overlaps between different composition structures and find that BFs belonging to different composition structures may partially overlap but some composition structures may in fact be subsets of other composition structures, e.g. {2,2} is a subset of {1,1,2}. Following this, we quantify the overlaps between composition structures and biologically meaningful BFs. Interestingly, we find that of the 9 composition structures (up to 5-input BFs), the NCFs are a subset of 6 composition structures.

Our final set of analyses are a set of 3 statistical tests to identify whether composed BFs are enriched within biological logics and if so, what are the factors that drive their enrichment. First, we find that the composed BFs are indeed enriched in our reference biological dataset in comparison to the space of all BFs. Then by computing the relative enrichment of a biologically meaningful sub-type in non-trivial composition structures (for instance, the relative enrichment of NCFs when considering BFs compatible with the composition structure {2,2}), we find that these sub-types are enriched, though the cause of its enrichment could be attributed either to the property of being biologically meaningful or to the property of belonging to the composition structure. To decide between these two possibilities, we compare the relative enrichments of biologically meaningful BFs which do not belong to the composed BFs to the relative enrichment of the composed BFs which do not belong to the biologically meaningful BFs. In a nutshell, these tests confirm that the property of being minimally complex in terms of the Boolean complexity or the average sensitivity, i.e., being either an RoF or a NCF, is most likely what drives the enrichment of composition structures.

## Appendix A: Properties of composed BFs

### Property A.1.

Given a composition structure {*t*_1_, *t*_2_,…, *t_r_*}, if *h* is a possible composed BF, then its complement 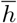 is also a possible composed BF.

*Proof*: *h* is a composed BF of the form *g*(*p*_1_, *p*_2_,…, *p_r_*), where each *p_i_* is a BF of *t_i_* inputs. There are 2^2*t*_1_^ 2^2*t*_2_^ … 2^2*t_r_*^ 2^2*^r^*^ combinations of *p*_1_, *p*_2_,…, *p_r_* and *g* which comprise the composed BFs *h*. Let us consider one such combination 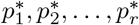 and *g*\*, that corresponds to a BF *h*\* from the space of all composed BFs. Now, there also exists another combination 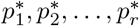 and 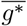 among the 2^2*t*_1_^ 2^2*t*_2_^ … 2^2*t_r_*^ 2^2*^r^*^ combinations, which corresponds to the BF 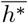. Hence, a function and its complement are both present in the composed BFs of any composition structure.

### Property A.2.

The composition structure {*k*} does not restrict the space of BFs.

*Proof*: The composition structure {*k*} corresponds to a *k*-input BF of the form *h* = *g*(*p*_1_(*x*_1_, *x*_2_,…, *x_k_*)). There are 2^2*^k^*^ BFs that can be assigned to *p*_1_, and 2^2^1^^ = 4 BFs that can be assigned to *g*. Among the 4 BFs that can be assigned to *g*, if we consider the *g*(*p*_1_) = *p*_1_, then *h* = *p*_1_(*x*_1_, *x*_2_,…, *x*_k_). It follows that any BF among the 2^2*^k^*^ possible *k*-input BFs can be assigned to *h*. Since *h* spans all the possible *k*-input BFs, the composition structure {*k*} cannot restrict the space of BFs.

## Appendix B: Complexes in Yeast

To perform the empirical analysis on *S. cerevisiae*, we first obtained a list of 617 macromolecular complexes in *S. cerevisiae* from the EBI Complex portal database [55]. Then we obtained the list of TFs in *S. cerevisiae* from the Yeastract database [70]. To do this, we obtained a list of 5195 verified genes from the SGD YeastMine database [71], and provided these genes as input to the Yeastract database. Additionally, we used the query “DNA binding evidence or expression evidence” in the Yeastract database. “DNA binding evidence” includes TF regulation verified by experiments such as EMSA, ChIP, ChIPchip and ChIP-seq, DNA footprinting, whereas “expression evidence” includes those interactions established by comparing gene expression in wild-type strains with mutant strains in which the gene encoding the TF is mutated. Expression evidence is obtained via northern blotting, RT-PCR, DNA microarrays and RNA-seq experiments [70]. The above query in the Yeastract database resulted in a list of 217 TFs that were used for further analysis. Note that 153 out of these 217 TFs show evidence for both DNA binding and effects on expression. Finally, among the 617 macromolecular complexes we selected only those complexes in which all the protein subunits correspond to TFs.

Using the EBI complex portal and the TFs from Yeastract database, we found that there are 17 heteromeric complexes among the 617 complexes in *S. cerevisiae* such that each of their protein subunits correspond to TFs (see Supplementary Table S3). Thereafter, we ascertained via manual curation of the literature associated with these 17 complexes that 15 of them act as TRs, 1 of them binds to DNA but whether it regulates gene expression is uncertain, and the remaining 1 acts as a transcriptional co-repressor. Among the 15 complexes that act as TRs, 9 complexes are formed by 2 proteins, 5 complexes are formed by 3 proteins and 1 complex is formed by 4 proteins. There are 30 unique TFs whose combination results in these 15 complexes. Additionally, we found that 11 out of these 15 complexes are such that their protein subunits show both DNA binding and expression evidence.

It is known from experiments that TFs belonging to the basic leucine zipper factors (bZIP) [57, 58] and basic helix-loop-helix factors (bHLH) [59, 60] classes typically bind DNA as dimers. To determine the classes of the 217 TFs in *S. cerevisiae,* we utilized the JASPAR database [61]. In *S. cerevisiae,* we found 10 TFs that belong to the bZIP class and 7 TFs that belong to the bHLH class (see Supplementary Table S4). Further, we find that all the 17 TFs in the bZIP or bHLH classes display evidence for both DNA binding and effects on expression.

## Appendix C: Reference biological dataset of 2687 Boolean functions from reconstructed models

Here, we utilized the empirical dataset compiled by Subbaroyan *et al.* [23] consisting of 2687 BFs extracted from 88 published discrete models of biological systems to quantify the abundance of composed BFs in biological networks. This dataset of BFs, available for download at https://github.com/asamallab/MCBF/tree/main/biological_dataset, was compiled using information on published models in online repositories Cell Collective [29] (https://cellcollective.org/), GINSIM [26] (http://ginsim.org/) or BioModels [27] (http://www.ebi.ac.uk/biomodels/) and by manually retrieving information from published literature. The compiled collection of 88 published models spans a diverse range of biological processes from various kingdoms of life. Though most of the 88 models are Boolean, some models are not Boolean but nevertheless discrete, implying that each node *may* have more than 2 states in its discrete logic model. For such models, BFs alone were compiled into the dataset, i.e., nodes which had a binary state and whose inputs were restricted to binary states were included in the 2687 BFs.

## Appendix D: Statistical Tests

### Enrichments and relative enrichments

For a given number of inputs *k*, let *T* be the set of composed BFs allowed by the composition structure {*t*_1_, *t*_2_,…, *t_r_*}, with *t*_1_ + *t*_2_ + … + *t_r_* = *k*. Let *f*_0_ be the fraction occupied by the set *T* among the set of all *k*-input BFs. This fraction *f*_0_ is equal to the probability of obtaining a BF belonging to the set *T* when drawing at random (uniformly) within all *k*-input BFs. Let *f*_1_ denote the fraction of BFs belonging to *T* among all *k*-input BFs that are present in the reference biological dataset. *T*’s “enrichment factor”, *E*, is equal to the fraction *f*_1_/*f*_0_. An enrichment factor *E* > 1 implies that the composition structure {*t*_1_, *t*_2_,…, *t_r_*} is enriched in the reference biological dataset, whereas *E* < 1 implies that the composition structure is depleted.

Consider now a refinement of the previous notion of enrichment to probe the roles of biologically meaningful types. Specifically, let *s* be one of the four types of biologically meaningful BFs (UFs, CFs, NCFs, or RoFs). Denote by *T_s_* the subset of BFs of type *s* in *T*. We then define the “relative enrichment” *E_R_* of type *s* in *T* as *E_R_* = *f*_*s*,1_/*f*_*s*,0_. In that ratio, *f*_*s*,0_ (respectively *f*_*s*,1_) is the fraction of BFs in *T* that furthermore belong to *T_s_* (respectively the fraction of BFs in the intersection of *T* and the reference biological dataset that furthermore belong to *T_s_*). A relative enrichment close to 1 means that an enrichment of *T_s_* is driven by an enrichment of *T* (i.e., by the composition structure) rather than by the biologically meaningful sub-type *s*. In contrast, a large relative enrichment suggests that sub-type *s* is driving enrichment of such BFs in the reference biological dataset even after accounting for enrichment of BFs compatible with a given composition structure.

### Associated p-values

We first describe the procedure that we employed to test the significance of an enrichment factor E for the set *T* of composed BFs in the composition structure {*t*_1_, *t*_2_,…, *t_r_* }. A similar statistical test was presented and implemented by Subbaroyan *et al.* [23]. First, we introduce the null hypothesis, denoted by *H*_0_, in which all the *k*-input BFs in the reference biological dataset are drawn from a random ensemble comprised of uniformly distributed *k*-input BFs. The p-value associated with rejecting this null hypothesis *H*_0_ is computed as follows. Recall that *f*_0_ is the probability of choosing a *k*-input BF belonging to the set *T* from the random ensemble. Define *M* to be the number of *k*-input BFs in the reference biological dataset. Then, we draw *M* BFs from all *k*-input BFs in the random ensemble, and compute the probability of getting *m* BFs that also belong to the set *T*. This probability is equivalent to getting *m* successes when tossing a biased coin *M* times, and is thus given by the binomial distribution 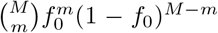. The *p*-value is then given by 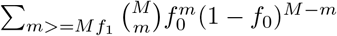. Here, *Mf*_1_ is the number of BFs belonging to the set *T* among all *k*-input BFs that are present in the reference biological dataset.

We perform a similar statistical test to determine whether a relative enrichment *E_R_* for a given *T* and *s* is statistically significant. In this case, the null hypothesis *H*_0_ hypothesizes that although there is a selection for *T* (as evident from a large value of *E*), the elements that are drawn within *T* have a uniform probability, that is the individual elements belonging to *T_s_* are not more probable than the other elements of *T*. In practice, we draw a sample of size M under *H*_0_ where as above *M* is the number of *k*-input BFs in the reference biological dataset. If this sample contains *M_T_* elements in *T* as in the reference biological dataset, the distribution of the number of elements in *T_s_* is then known. Specifically, the probability to have m elements in *T_s_* is 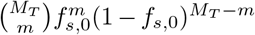 where *f*_*s*,0_ is the ratio of sizes of *T_s_* and *T* in the random ensemble. The *p*-value associated with rejecting *H*_0_ is then just the sum of all such probabilities under the condition that *m* is larger or equal to the number of *T_s_* elements in the reference biological dataset.

## ACKNOWLEDGMENTS

We are thankful to Satyanarayan Rao for discussions and Donghoon Lee for sharing the list of active enhancers from STARRPeaker. A. Samal acknowledges research support from the Department of Atomic Energy (DAE), India and Max Planck Society, Germany (through a Max Planck Partner Group in Mathematical Biology). IPS2 benefits from the support of Saclay Plant Sciences-SPS (ANR-17-EUR-0007).

## DATA AVAILABILITY

All datasets and programs needed to reproduce the results of this study are available from the associated GitHub repository: https://github.com/asamallab/CoSt

## References

[1] K. Chen and N. Rajewsky. The evolution of gene regulation by transcription factors and microRNAs. Nature Reviews Genetics, 8(2):93–103, 2007.

[2] R. Milo, S. Shen-Orr, S. Itzkovitz, N. Kashtan, D. Chklovskii, and U. Alon. Network Motifs: Simple Building Blocks of Complex Networks. Science, 298(5594):824–827, 2002.

[3] A. L. Barabási and Z. N. Oltvai. Network biology: understanding the cell’s functional organization. Nature Reviews Genetics, 5(2):101–113, 2004.

[4] S. Bornholdt. Less is more in modeling large genetic networks. Science, 310(5747):449–451, 2005.

[5] U. Alon. An Introduction to Systems Biology: Design Principles of Biological Circuits. Chapman and Hall/CRC., 2006.

[6] S. A. Kauffman. Metabolic stability and epigenesis in randomly constructed genetic nets. Journal of Theoretical Biology, 22(3):437–467, 1969.

[7] S. A. Kauffman. Homeostasis and differentiation in random genetic control networks. Nature, 224(5215):177–178, 1969.

[8] R. Thomas. Boolean formalization of genetic control circuits. Journal of Theoretical Biology, 42(3):563–585, 1973.

[9] R. Thomas. Kinetic logic: a Boolean approach to the analysis of complex regulatory systems, Proceedings of the EMBO course “Formal analysis of genetic regulation”, held in Brussels, September 6–16, 1977, Lecture notes in Biomathematics, volume 29. Springer, 1979.

[10] S. A. Kauffman, C. Peterson, B. Samuelsson, and C. Troein. Random Boolean network models and the yeast transcriptional network. Proceedings of the National Academy of Sciences, 100(25):14796–14799, 2003.

[11] M. Davidich and S. Bornholdt. The transition from differential equations to Boolean networks: a case study in simplifying a regulatory network model. Journal of Theoretical Biology, 255(3):269–277, 2008.

[12] J. Saez-Rodriguez, L. Simeoni, J. A. Lindquist, R. Hemenway, U. Bommhardt, et al. A logical model provides insights into T cell receptor signaling. PLoS Computational Biology, 3(8):e163, 2007.

[13] A. Samal and S. Jain. The regulatory network of E. coli metabolism as a Boolean dynamical system exhibits both homeostasis and flexibility of response. BMC Systems Biology, 2(21):1–18, 2008.

[14] A. L. Bauer, T. L. Jackson, Y. Jiang, and T. Rohlf. Receptor cross-talk in angiogenesis: mapping environmental cues to cell phenotype using a stochastic, Boolean signaling network model. Journal of Theoretical Biology, 264(3):838–846, 2010.

[15] I. Shmulevich and S. A. Kauffman. Activities and sensitivities in Boolean network models. Physical Review Letters, 93(4):48701, 2004.

[16] B. Drossel, T. Mihaljev, and F. Greil. Number and length of attractors in a critical Kauffman model with connectivity one. Physical Review Letters, 94(8):088701, 2005.

[17] K. Klemm and S. Bornholdt. Stable and unstable attractors in Boolean networks. Physical Review E, 72(5):055101, 2005.

[18] B. Ø Palsson. Systems Biology: Properties of Reconstructed Networks. Cambridge University Press, 2006.

[19] M. Nykter, N. D. Price, M. Aldana, S. A Ramsey, S. A. Kauffman, et al. Gene expression dynamics in the macrophage exhibit criticality. Proceedings of the National Academy of Sciences, 105(6):1897–1900, 2008.

[20] E. Balleza, E. R. Alvarez-Buylla, A. Chaos, S. Kauffman, I. Shmulevich, and M. Aldana. Critical dynamics in genetic regulatory networks: examples from four kingdoms. PLoS One, 3(6):e2456, 2008.

[21] S. Chowdhury, J. Lloyd-Price, O. Smolander, W. C. Baici, T. R. Hughes, et al. Information propagation within the genetic network of Saccharomyces cerevisiae. BMC Systems Biology, 4(1):1–10, 2010.

[22] B. C. Daniels, H. Kim, D. Moore, S. Zhou, H. B. Smith,B. Karas, S. A. Kauffman, and S. I. Walker. Criticality Distinguishes the Ensemble of Biological Regulatory Networks. Physical Review Letters, 121(13):138102, 2018.

[23] A. Subbaroyan, O. C. Martin, and A. Samal. Minimum complexity drives regulatory logic in boolean models of living systems. PNAS Nexus, 1:pgac017, 2022.

[24] L. Mendoza, D. Thieffry, and E. R. Alvarez-Buylla. Genetic control of flower morphogenesis in Arabidopsis thaliana: a logical analysis. Bioinformatics, 15(7):593–606, 1999.

[25] R. Albert and H. G. Othmer. The topology of the regulatory interactions predicts the expression pattern of the segment polarity genes in *Drosophila melanogaster*. Journal of Theoretical Biology, 223(1):1–18, 2003.

[26] A. G. Gonzalez, A. Naldi, L. Sanchez, D. Thieffry, and C. Chaouiya. GINsim: A software suite for the qualitative modelling, simulation and analysis of regulatory networks. BioSystems, 84(2):91–100, 2006.

[27] C. Li, M. Donizelli, N. Rodriguez, H. Dharuri, L. Endler, et al. BioModels Database: An enhanced, curated and annotated resource for published quantitative kinetic models. BMC Systems Biology, 4(1):1–14, 2010.

[28] A. Saadatpour, R. S. Wang, A. Liao, X. Liu, T. P. Loughran, I. Albert, and R. Albert. Dynamical and Structural Analysis of a T Cell Survival Network Identifies Novel Candidate Therapeutic Targets for Large Granular Lymphocyte Leukemia. PLOS Computational Biology, 7(11):e1002267, 2011.

[29] T. Helikar, B. Kowal, S. McClenathan, M. Bruckner, T. Rowley, A. Madrahimov, B. Wicks, M. Shrestha, K. Limbu, and J. A. Rogers. The Cell Collective: Toward an open and collaborative approach to systems biology. BMC Systems Biology, 6(1):1–14, 2012.

[30] A. Méndez and L. Mendoza. A Network Model to Describe the Terminal Differentiation of B Cells. PLOS Computational Biology, 12(1):e1004696, 2016.

[31] E. Guberman, H. Sherief, and E. R. Regan. Boolean model of anchorage dependence and contact inhibition points to coordinated inhibition but semi-independent induction of proliferation and migration. Computational and Structural Biotechnology Journal, 18:2145–2165, 2020.

[32] J. Aracena. Maximum Number of Fixed Points in Regulatory Boolean Networks. Bulletin of Mathematical Biology, 70(5):1398, 2008.

[33] S. A. Kauffman. The origins of order: self-organization and selection in evolution. Oxford University Press, New York, 1993.

[34] A. S. Jarrah, B. Raposa, and R. Laubenbacher. Nested Canalyzing, Unate Cascade, and Polynomial Functions. Physica D: Nonlinear Phenomena, 233(2):167–174, 2007.

[35] C. Kadelka, J. Kuipers, and R. Laubenbacher. The influence of canalization on the robustness of Boolean networks. Physica D: Nonlinear Phenomena, 353:39–47, 2017.

[36] B. Schwanhäusser, D. Busse, N. Li, G. Dittmar, J. Schuchhardt, J. Wolf, W. Chen, and M. Selbach. Global quantification of mammalian gene expression control. Nature, 473(7347):337–342, 2011.

[37] R. Hannam, R. Kühn, and A. Annibale. Percolation in bipartite Boolean networks and its role in sustaining life. Journal of Physics A: Mathematical and Theoretical, 52(33):334002, 2019.

[38] M. Flöttmann, F. Krause, E. Klipp, and M. Krantz. Reaction-contingency based bipartite Boolean modelling. BMC Systems Biology, 7(1):1–12, 2013.

[39] T. Mori, M. Flöttmann, M. Krantz, T. Akutsu, and E. Klipp. Stochastic simulation of Boolean rxncon models: towards quantitative analysis of large signaling networks. BMC Systems Biology, 9(1):1–9, 2015.

[40] G. Torrisi, R. Kühn, and A. Annibale. Percolation on the gene regulatory network. Journal of Statistical Mechanics: Theory and Experiment, 2020(8):083501, 2020.

[41] T. Fink and R. Hannam. Boolean composition restricts biological logics. arXiv preprint arXiv:2109.12551, 2021.

[42] I. Shmulevich, H. Lähdesmäki, E. R. Dougherty, J. Astola, and W. Zhang. The role of certain post classes in Boolean network models of genetic networks. Proceedings of the National Academy of Sciences, 100(19):10734–10739, 2003.

[43] D. Shlyueva, G. Stampfel, and A. Stark. Transcriptional enhancers: from properties to genome-wide predictions. Nature Reviews Genetics, 15(4):272–286, 2014.

[44] F. Reiter, S. Wienerroither, and A. Stark. Combinatorial function of transcription factors and cofactors. Current opinion in genetics & development, 43:73–81, 2017.

[45] R. Thomas. Regulatory networks seen as asynchronous automata: a logical description. Journal of Theoretical Biology, 153(1):1–23, 1991.

[46] J. Feldman. A catalog of Boolean concepts. Journal of Mathematical Psychology, 47(1):75–89, 2003.

[47] Z. Szallasi and S. Liang. Modeling the normal and neoplastic cell cycle with ‘realistic Boolean genetic networks’: Their application for understanding carcinogenesis and assessing therapeutic strategies. In Pacific Symposium on Biocomputing, volume 3, pages 66–76. Citeseer, 1998.

[48] M. C. Golumbic and V. Gurvich. Read-once functions, page 448–486. Encyclopedia of Mathematics and its Applications. Cambridge University Press, 2011.

[49] H. Rottensteiner, A. J. Kal, B. Hamilton, H. Ruis, and H. F. Tabak. A heterodimer of the Zn2Cys6 transcription factors Pip2p and Oaf1p controls induction of genes encoding peroxisomal proteins in Saccharomyces cerevisiae. European Journal of Biochemistry, 247(3):776–783, 1997.

[50] L. Fernandes, C. Rodrigues-Pousada, and K. Struhl. Yap, a novel family of eight bZIP proteins in *Saccharomyces cerevisiae* with distinct biological functions. Molecular and Cellular Biology, 17(12):6982–6993, 1997.

[51] C. Wolberger. Multiprotein-DNA Complexes in Transcriptional Regulation. Annual Review of Biophysics and Biomolecular Structure, 28(1):29–56, 1999.

[52] T. Vernoux, G. Brunoud, E. Farcot, V. Morin, H. Van den Daele, et al. The auxin signalling network translates dynamic input into robust patterning at the shoot apex. Molecular Systems Biology, 7(1):508, 2011.

[53] A. P. W. Funnell and M. Crossley. Homo-and Heterodimerization in Transcriptional Regulation. In Protein Dimerization and Oligomerization in Biology, volume 747, pages 105–121. Springer New York, New York, NY, 2012.

[54] T. J. Guilfoyle and G. Hagen. Auxin response factors. Current opinion in plant biology, 10(5):453–460, 2007.

[55] B.H.M. Meldal, L. Perfetto, C. Combe, T. Lubiana, J. V. Ferreira Cavalcante, H. Bye-A-Jee, A. Waagmeester, N. Del-Toro, A. Shrivastava, E. Barrera, et al. Complex Portal 2022: new curation frontiers. Nucleic Acids Research, 50(D1):D578–D586, 2022.

[56] S. A. Lambert, A. Jolma, L. F. Campitelli, P. K. Das, Y. Yin, et al. The human transcription factors. Cell, 172(4):650–665, 2018.

[57] J. A. Rodríiguez-Martínez, A. W. Reinke, D. Bhimsaria, A. E. Keating, and A. Z. Ansari. Combinatorial bZIP dimers display complex DNA-binding specificity landscapes. eLife, 6:e19272, 2017.

[58] W. Dröge-Laser, B. L. Snoek, B. Snel, and C. Weiste. The Arabidopsis bZIP transcription factor family — an update. Current Opinion in Plant Biology, 45:36–49, 2018.

[59] S. Jones. An overview of the basic helix-loop-helix proteins. Genome Biology, 5(226):6, 2004.

[60] L. Chen and J. M. Lopes. Multiple bHLH proteins regulate CIT2 expression in Saccharomyces cerevisiae. Yeast, 27(6):345–359, 2010.

[61] J. A. Castro-Mondragon, R. Riudavets-Puig, I. Rauluseviciute, R. Berhanu Lemma, L. Turchi, et al. JASPAR 2022: the 9th release of the open-access database of transcription factor binding profiles. Nucleic Acids Research, 50(D1):D165–D173, 2021.

[62] F. Spitz and E. E. Furlong. Transcription factors: from enhancer binding to developmental control. Nature Reviews Genetics, 13(9):613–626, 2012.

[63] A. Sandelin, P. Carninci, B. Lenhard, J. Ponjavic, Y. Hayashizaki, and D. A. Hume. Mammalian RNA polymerase II core promoters: insights from genome-wide studies. Nature Reviews Genetics, 8(6):424–436, 2007.

[64] E. M. Blackwood and J. T. Kadonaga. Going the distance: a current view of enhancer action. Science, 281(5373):60–63, 1998.

[65] S. Rao, K. Ahmad, and S. Ramachandran. Cooperative binding between distant transcription factors is a hallmark of active enhancers. Molecular Cell, 81(8):1651–1665.e4, 2021.

[66] ENCODE Project Consortium et al. An integrated encyclopedia of DNA elements in the human genome. Nature, 489(7414):57, 2012.

[67] D. Lee, M. Shi, J. Moran, M. Wall, J. Zhang, et al. STARRPeaker: uniform processing and accurate identification of STARR-seq active regions. Genome biology, 21(1):1–24, 2020.

[68] A. Lex, N. Gehlenborg, H. Strobelt, R. Vuillemot, and H. Pfister. UpSet: Visualization of Intersecting Sets. IEEE transactions on visualization and computer graphics, 20(12):1983–1992, 2014.

[69] S. Nikolajewa, M. Friedel, and T. Wilhelm. Boolean networks with biologically relevant rules show ordered behavior. BioSystems, 90(1):40–47, 2007.

[70] M. C. Teixeira, P. T. Monteiro, M. Palma, C. Costa, C. P. Godinho, et al. Yeastract: an upgraded database for the analysis of transcription regulatory networks in Saccharomyces cerevisiae. Nucleic Acids Research, 46(D1):D348–D353, 2018.

[71] R. Balakrishnan, J. Park, K. Karra, B. C. Hitz, G. Binkley, E. L. Hong, J. Sullivan, G. Micklem, and J. M. Cherry. Yeastmine—an integrated data warehouse for *Saccharomyces cerevisiae* data as a multipurpose toolkit. Database, 2012, 2012.

