## Supplementary Table for "Composition structures and biologically meaningful logics: plausibility and relevance in bipartite models of gene regulation"

for

TABLE S1: **A list of 169 complexes from the set of 1325 complexes in *H. sapiens* such that all the protein subunits of these complexes are transcription factors.** The 1325 complexes in *H. sapiens* were obtained from the EBI Complex Portal database. The list of human TFs was obtained from `http://humantfs.ccbr.utoronto.ca/`. 'Complex ID' is the identifier of the complex as given in the EBI Complex Portal database. 'Complex name' is the name of the complex as given in the EBI Complex Portal database. 'Size of the complex' is the number of protein subunits the complex is constituted of. 'Uniprot ID of protein subunits' gives the Uniprot IDs of the subunits in a complex and their stoichiometric coefficients as integers within brackets. 'Is a TR' column is 'yes' if the complex acts as a transcriptional regulator (TR) based on manual literature curation.

| Complex ID | Complex name | Size of the complex | UniProt ID of protein subunits | Is a TF |
|---|---|---|---|---|
| CPX-6405 | bZIP transcription factor complex, ATF1-CREB1 | 2 | P16220(1)<br>P18846(1) | yes |
| CPX-6414 | bZIP transcription factor complex, ATF2-BATF3 | 2 | P15336(1)<br>Q9NR55(1) | yes |
| CPX-6416 | bZIP transcription factor complex, ATF2-FOS | 2 | P01100(1)<br>P15336(1) | yes |
| CPX-6420 | bZIP transcription factor complex, ATF2-JUN | 2 | P05412(1)<br>P15336(1) | yes |
| CPX-6421 | bZIP transcription factor complex, ATF2-JUNB | 2 | P15336(1)<br>P17275(1) | yes |
| CPX-6467 | bZIP transcription factor complex, ATF3-BATF | 2 | P18847(1)<br>Q16520(1) | yes |
| CPX-6468 | bZIP transcription factor complex, ATF3-BATF3 | 2 | P18847(1)<br>Q9NR55(1) | yes |
| CPX-6469 | bZIP transcription factor complex, ATF3-CEBPA | 2 | P18847(1)<br>P49715(1) | yes |
| CPX-6471 | bZIP transcription factor complex, ATF3-CEBPG | 2 | P17676(1)<br>P18847(1) | yes |
| CPX-6477 | bZIP transcription factor complex, ATF3-FOS | 2 | P01100(1)<br>P18847(1) | yes |
| CPX-6478 | bZIP transcription factor complex, ATF3-FOSL1 | 2 | P15407(1)<br>P18847(1) | yes |
| CPX-6474 | bZIP transcription factor complex, ATF3-JUN | 2 | P05412(1)<br>P18847(1) | yes |
| CPX-6476 | bZIP transcription factor complex, ATF3-JUNB | 2 | P17275(1)<br>P18847(1) | yes |
| CPX-6523 | bZIP transcription factor complex, ATF4-BATF2 | 2 | P18848(1)<br>Q8N1L9(1) | yes |
| CPX-6524 | bZIP transcription factor complex, ATF4-BATF3 | 2 | P18848(1)<br>Q9NR55(1) | yes |

| | | | | |
|---|---|---|---|---|
| CPX-6525 | bZIP transcription factor complex, ATF4-CEBPA | 2 | P18848(1)<br>P49715(1) | yes |
| CPX-6527 | bZIP transcription factor complex, ATF4-CEBPG | 2 | P18848(1)<br>P53567(1) | yes |
| CPX-6563 | bZIP transcription factor complex, ATF4-JUNB | 2 | P17275(1)<br>P18848(1) | yes |
| CPX-6567 | bZIP transcription factor complex, ATF4-MAFB | 2 | P18848(1)<br>Q9Y5Q3(1) | yes |
| CPX-6585 | bZIP transcription factor complex, ATF5-BATF | 2 | Q16520(1)<br>Q9Y2D1(1) | yes |
| CPX-6586 | bZIP transcription factor complex, ATF5-CEBPA | 2 | P49715(1)<br>Q9Y2D1(1) | yes |
| CPX-6589 | bZIP transcription factor complex, ATF5-CEBPE | 2 | Q15744(1)<br>Q9Y2D1(1) | yes |
| CPX-6588 | bZIP transcription factor complex, ATF5-CEBPG | 2 | P53567(1)<br>Q9Y2D1(1) | yes |
| CPX-7006 | bZIP transcription factor complex, BATF-CEBPA | 2 | P49715(1)<br>Q16520(1) | yes |
| CPX-7010 | bZIP transcription factor complex, BATF-CEBPE | 2 | Q15744(1)<br>Q16520(1) | yes |
| CPX-7008 | bZIP transcription factor complex, BATF-CEBPG | 2 | P53567(1)<br>Q16520(1) | yes |
| CPX-7011 | bZIP transcription factor complex, BATF-HLF | 2 | Q16520(1)<br>Q16534(1) | yes |
| CPX-7003 | bZIP transcription factor complex, BATF-JUNB | 2 | P17275(1)<br>Q16520(1) | yes |
| CPX-7017 | bZIP transcription factor complex, BATF-NFIL3 | 2 | Q16520(1)<br>Q16649(1) | yes |
| CPX-7063 | bZIP transcription factor complex, BATF2-JUN | 2 | P05412(1)<br>Q8N1L9(1) | yes |
| CPX-7061 | bZIP transcription factor complex, BATF2-JUNB | 2 | P17275(1)<br>Q8N1L9(1) | yes |
| CPX-7095 | bZIP transcription factor complex, BATF3-CEBPA | 2 | P49715(1)<br>Q9NR55(1) | yes |
| CPX-7097 | bZIP transcription factor complex, BATF3-CEBPG | 2 | P53567(1)<br>Q9NR55(1) | yes |
| CPX-7100 | bZIP transcription factor complex, BATF3-JUN | 2 | P05412(1)<br>Q9NR55(1) | yes |
| CPX-7101 | bZIP transcription factor complex, BATF3-JUNB | 2 | P17275(1)<br>Q9NR55(1) | yes |
| CPX-486 | bZIP transcription factor complex, FOS-JUN | 2 | P01100(1)<br>P05412(1) | yes |
| CPX-504 | c-Myb-C/EBPbeta complex | 2 | P10242(1)<br>P17676(2) | yes |

| | | | | |
|---|---|---|---|---|
| CPX-1956 | CCAAT-binding factor complex | 3 | P23511(1)<br>P25208(1)<br>Q13952(1) | yes |
| CPX-3229 | CLOCK-BMAL1 transcription complex | 2 | O00327(1)<br>O15516(1) | yes |
| CPX-3230 | CLOCK-BMAL2 transcription complex | 2 | O15516(1)<br>Q8WYA1(1) | yes |
| CPX-5156 | ERalpha-NCOA2 activated estrogen receptor complex | 2 | P03372(2)<br>Q15596(2) | yes |
| CPX-1123 | FOXO3-MYC complex | 2 | O43524(0)<br>P01106(0) | yes |
| CPX-6016 | ISGF3 complex | 3 | P42224(0)<br>P52630(0)<br>Q00978(0) | yes |
| CPX-5834 | NF-kappaB DNA-binding transcription factor complex, p65/c-Rel | 2 | Q04206(1)<br>Q04864(1) | yes |
| CPX-517 | PXR-NCOA1 activated nuclear receptor complex | 2 | O75469(2)<br>Q15788(2) | yes |
| CPX-496 | RXRalpha-PXR nuclear receptor complex | 2 | O75469(2)<br>P19793(2) | yes |
| CPX-508 | RXRalpha-RARalpha retinoic acid receptor complex | 2 | P10276(1)<br>P19793(1) | yes |
| CPX-654 | RXRalpha-TRbeta nuclear hormone receptor complex | 2 | P10828(1)<br>P19793(1) | yes |
| CPX-54 | SMAD1-SMAD4 complex | 2 | Q13485(1)<br>Q15797(2) | yes |
| CPX-3252 | SMAD3-SMAD4 complex | 2 | P84022(2)<br>Q13485(1) | yes |
| CPX-6041 | STAT1/STAT3 complex | 2 | P40763(1)<br>P42224(1) | yes |
| CPX-6042 | STAT1/STAT4 complex | 2 | P42224(1)<br>Q14765(1) | yes |
| CPX-6043 | STAT3/STAT5A complex | 2 | P40763(1)<br>P42229(1) | yes |
| CPX-6044 | STAT3/STAT5B complex | 2 | P40763(1)<br>P51692(1) | yes |
| CPX-91 | Transcriptional activator Myc-Max complex | 2 | P01106(1)<br>P61244(1) | yes |
| CPX-104 | Transcriptional repressor Mad-Max complex | 2 | P61244(1)<br>Q05195(1) | yes |
| CPX-3079 | USF1-USF2 upstream stimulatory factor complex | 2 | P22415(1)<br>Q15853(1) | yes |
| CPX-6419 | bZIP transcription factor complex, ATF2-JDP2 | 2 | P15336(1)<br>Q8WYK2(1) | yes |

| | | | | |
|---|---|---|---|---|
| CPX-6422 | bZIP transcription factor complex, ATF2-JUND | 2 | P15336(1) P17535(1) | yes |
| CPX-6595 | bZIP transcription factor complex, ATF6-ATF6B | 2 | P18850(1) Q99941(1) | yes |
| CPX-6600 | bZIP transcription factor complex, ATF6B-XBP1 | 2 | P17861(1) Q99941(1) | yes |
| CPX-2500 | bZIP transcription factor complex, BACH1-MAF | 2 | O14867(1) O75444(1) | yes |
| CPX-7165 | bZIP transcription factor complex, BACH1-MAFF | 2 | O14867(1) Q9ULX9(1) | yes |
| CPX-2872 | bZIP transcription factor complex, BACH1-MAFG | 2 | O14867(1) O15525(1) | yes |
| CPX-2493 | bZIP transcription factor complex, BACH1-MAFK | 2 | O14867(1) O60675(1) | yes |
| CPX-2483 | bZIP transcription factor complex, BACH2-MAF | 2 | O75444(1) Q9BYV9(1) | yes |
| CPX-2484 | bZIP transcription factor complex, BACH2-MAFF | 2 | Q9BYV9(1) Q9ULX9(1) | yes |
| CPX-2482 | bZIP transcription factor complex, BACH2-MAFK | 2 | O60675(1) Q9BYV9(1) | yes |
| CPX-7005 | bZIP transcription factor complex, BATF-JUN | 2 | P05412(1) Q16520(1) | yes |
| CPX-7102 | bZIP transcription factor complex, BATF3-JUND | 2 | P17535(1) Q9NR55(1) | yes |
| CPX-509 | bZIP transcription factor complex, CEBPA-CEBPB | 2 | P17676(1) P49715(1) | yes |
| CPX-1971 | E2F1-DP1 transcription factor complex | 2 | Q01094(1) Q14186(1) | yes |
| CPX-1972 | E2F2-DP1 transcription factor complex | 2 | Q14186(1) Q14209(1) | yes |
| CPX-711 | PPARgamma-NCOA1 activated nuclear receptor complex | 2 | P37231(1) Q15788(1) | yes |
| CPX-702 | PPARgamma-NCOA2 activated nuclear receptor complex | 2 | P37231(1) Q15596(1) | yes |
| CPX-525 | RARalpha-NCOA1 activated retinoic acid receptor complex | 2 | P10276(2) Q15788(2) | yes |
| CPX-666 | RARalpha-NCOA2 activated retinoic acid receptor complex | 2 | P10276(2) Q15596(2) | yes |
| CPX-632 | RXRalpha-LXRalpha nuclear hormone receptor complex | 2 | P19793(1) Q13133(1) | yes |
| CPX-678 | RXRalpha-LXRbeta nuclear hormone receptor complex | 2 | P19793(1) P55055(1) | yes |
| CPX-513 | RXRalpha-NCOA2 activated retinoic acid receptor complex | 2 | P19793(2) Q15596(2) | yes |

| | | | | |
|---|---|---|---|---|
| CPX-816 | RXRalpha-RARalpha-NCOA2 retinoic acid receptor complex | 3 | P10276(1)<br>P19793(1)<br>Q15596(0) | yes |
| CPX-631 | RXRalpha-VDR nuclear hormone receptor complex | 2 | P11473(1)<br>P19793(1) | yes |
| CPX-716 | RXRbeta-LXRalpha nuclear hormone receptor complex | 2 | P28702(1)<br>Q13133(1) | yes |
| CPX-652 | RXRbeta-LXRbeta nuclear hormone receptor complex | 2 | P28702(1)<br>P55055(1) | yes |
| CPX-871 | RXRbeta-VDR nuclear hormone receptor complex | 2 | P11473(1)<br>P28702(1) | yes |
| CPX-6062 | SMAD3-TTF-1 complex | 2 | P43699(0)<br>P84022(0) | yes |
| CPX-2497 | bZIP transcription factor complex, BACH1-MAFB | 2 | O14867(1)<br>Q9Y5Q3(1) | uncertain |
| CPX-6407 | bZIP transcription factor complex, ATF2-ATF3 | 2 | P15336(1)<br>P18847(1) | uncertain |
| CPX-6408 | bZIP transcription factor complex, ATF2-ATF4 | 2 | P15336(1)<br>P18848(1) | uncertain |
| CPX-6415 | bZIP transcription factor complex, ATF2-DDIT3 | 2 | P15336(1)<br>P35638(1) | uncertain |
| CPX-6417 | bZIP transcription factor complex, ATF2-FOSL1 | 2 | P15336(1)<br>P15407(1) | uncertain |
| CPX-6385 | bZIP transcription factor complex, ATF3-ATF4 | 2 | P18847(1)<br>P18848(1) | uncertain |
| CPX-6472 | bZIP transcription factor complex, ATF3-CEBPE | 2 | P18847(1)<br>Q15744(1) | uncertain |
| CPX-6473 | bZIP transcription factor complex, ATF3-DDIT3 | 2 | P18847(1)<br>P35638(1) | uncertain |
| CPX-6542 | bZIP transcription factor complex, ATF4-CREBZF | 2 | P18848(1)<br>Q9NS37(1) | uncertain |
| CPX-6543 | bZIP transcription factor complex, ATF4-DDIT3 | 2 | P18848(1)<br>P35638(1) | uncertain |
| CPX-6564 | bZIP transcription factor complex, ATF4-FOS | 2 | P01100(1)<br>P18848(1) | uncertain |
| CPX-6565 | bZIP transcription factor complex, ATF4-FOSL1 | 2 | P15407(1)<br>P18848(1) | uncertain |
| CPX-6562 | bZIP transcription factor complex, ATF4-JUN | 2 | P05412(1)<br>P18848(1) | uncertain |
| CPX-6597 | bZIP transcription factor complex, ATF6-XBP1 | 2 | P17861(1)<br>P18850(1) | uncertain |
| CPX-2485 | bZIP transcription factor complex, BACH2-MAFG | 2 | O15525(1)<br>Q9BYV9(1) | uncertain |

| | | | | |
|---|---|---|---|---|
| CPX-7014 | bZIP transcription factor complex, BATF-DBP | 2 | Q10586(1) Q16520(1) | uncertain |
| CPX-7004 | bZIP transcription factor complex, BATF-DDIT3 | 2 | P35638(1) Q16520(1) | uncertain |
| CPX-7108 | bZIP transcription factor complex, BATF3-DBP | 2 | Q10586(1) Q9NR55(1) | uncertain |
| CPX-7106 | bZIP transcription factor complex, BATF3-DDIT3 | 2 | P35638(1) Q9NR55(1) | uncertain |
| CPX-69 | bZIP transcription factor complex, CEBPA-DDIT3 | 2 | P35638(1) P49715(1) | uncertain |
| CPX-70 | bZIP transcription factor complex, CEBPB-DDIT3 | 2 | P17676(1) P35638(1) | uncertain |
| CPX-6047 | STAT2/STAT6 complex | 2 | P42226(1) P52630(1) | uncertain |
| CPX-6046 | STAT3/STAT4 complex | 2 | P40763(1) Q14765(1) | uncertain |
| CPX-6045 | STAT5A/STAT5B complex | 2 | P42229(1) P51692(1) | uncertain |
| CPX-480 | AP-1 transcription factor complex FOS-JUN-NFATC2 | 3 | P01100(1) P05412(1) Q13469(1) | uncertain |
| CPX-9 | bZIP transcription factor complex, ATF1-ATF4 | 2 | P18846(1) P18848(1) | uncertain |
| CPX-6402 | bZIP transcription factor complex, ATF1-BACH1 | 2 | O14867(1) P18846(1) | uncertain |
| CPX-6404 | bZIP transcription factor complex, ATF1-NFIL3 | 2 | P18846(1) Q16649(1) | uncertain |
| CPX-6409 | bZIP transcription factor complex, ATF2-ATF7 | 2 | P15336(1) P17544(1) | uncertain |
| CPX-6412 | bZIP transcription factor complex, ATF2-BACH1 | 2 | O14867(1) P15336(1) | uncertain |
| CPX-6413 | bZIP transcription factor complex, ATF2-BATF | 2 | P15336(1) Q16520(1) | uncertain |
| CPX-6418 | bZIP transcription factor complex, ATF2-FOSL2 | 2 | P15336(1) P15408(1) | uncertain |
| CPX-6466 | bZIP transcription factor complex, ATF3-ATF7 | 2 | P17544(1) P18847(1) | uncertain |
| CPX-6470 | bZIP transcription factor complex, ATF3-CEBPB | 2 | P17676(1) P18847(1) | uncertain |
| CPX-6479 | bZIP transcription factor complex, ATF3-FOSL2 | 2 | P15408(1) P18847(1) | uncertain |
| CPX-6480 | bZIP transcription factor complex, ATF3-MAFF | 2 | P18847(1) Q9ULX9(1) | uncertain |

| | | | | |
|---|---|---|---|---|
| CPX-6481 | bZIP transcription factor complex, ATF3-MAFG | 2 | O15525(1) P18847(1) | uncertain |
| CPX-6522 | bZIP transcription factor complex, ATF4-BATF | 2 | P18848(1) Q16520(1) | uncertain |
| CPX-6526 | bZIP transcription factor complex, ATF4-CEBPB | 2 | P17676(1) P18848(1) | uncertain |
| CPX-6528 | bZIP transcription factor complex, ATF4-CEBPD | 2 | P18848(1) P49716(1) | uncertain |
| CPX-6529 | bZIP transcription factor complex, ATF4-CEBPE | 2 | P18848(1) Q15744(1) | uncertain |
| CPX-8 | bZIP transcription factor complex, ATF4-CREB1 | 2 | P16220(1) P18848(1) | uncertain |
| CPX-6541 | bZIP transcription factor complex, ATF4-CREB3 | 2 | O43889(1) P18848(1) | uncertain |
| CPX-6566 | bZIP transcription factor complex, ATF4-MAF | 2 | O75444(1) P18848(1) | uncertain |
| CPX-6568 | bZIP transcription factor complex, ATF4-NFE2 | 2 | P18848(1) Q16621(1) | uncertain |
| CPX-6570 | bZIP transcription factor complex, ATF4-NFE2L2 | 2 | P18848(1) Q16236(1) | uncertain |
| CPX-6572 | bZIP transcription factor complex, ATF4-NFE2L3 | 2 | P18848(1) Q9Y4A8(1) | uncertain |
| CPX-6601 | bZIP transcription factor complex, ATF6B-CREBZF | 2 | Q99941(1) Q9NS37(1) | uncertain |
| CPX-6781 | bZIP transcription factor complex, ATF7-BACH1 | 2 | O14867(1) P17544(1) | uncertain |
| CPX-6782 | bZIP transcription factor complex, ATF7-CEBPG | 2 | P17544(1) P53567(1) | uncertain |
| CPX-6784 | bZIP transcription factor complex, ATF7-DDIT3 | 2 | P17544(1) P35638(1) | uncertain |
| CPX-6783 | bZIP transcription factor complex, ATF7-FOS | 2 | P01100(1) P17544(1) | uncertain |
| CPX-6785 | bZIP transcription factor complex, ATF7-FOSL2 | 2 | P15408(1) P17544(1) | uncertain |
| CPX-6786 | bZIP transcription factor complex, ATF7-JUN | 2 | P05412(1) P17544(1) | uncertain |
| CPX-6787 | bZIP transcription factor complex, ATF7-JUNB | 2 | P17275(1) P17544(1) | uncertain |
| CPX-6788 | bZIP transcription factor complex, ATF7-JUND | 2 | P17535(1) P17544(1) | uncertain |
| CPX-6789 | bZIP transcription factor complex, ATF7-NFE2 | 2 | P17544(1) Q16621(1) | uncertain |
| CPX-7012 | bZIP transcription factor complex, BACH1-BATF | 2 | O14867(1) Q16520(1) | uncertain |

| | | | | |
|---|---|---|---|---|
| CPX-2494 | bZIP transcription factor complex, BACH1-CREB1 | 2 | O14867(1)<br>P16220(1) | uncertain |
| CPX-2496 | bZIP transcription factor complex, BACH1-DDIT3 | 2 | O14867(1)<br>P35638(1) | uncertain |
| CPX-2491 | bZIP transcription factor complex, BACH1-FOS | 2 | O14867(1)<br>P01100(1) | uncertain |
| CPX-7093 | bZIP transcription factor complex, BACH2-BATF3 | 2 | Q9BYV9(1)<br>Q9NR55(1) | uncertain |
| CPX-2479 | bZIP transcription factor complex, BACH2-MAFB | 2 | Q9BYV9(1)<br>Q9Y5Q3(1) | uncertain |
| CPX-2471 | bZIP transcription factor complex, BACH2-NFE2L3 | 2 | Q9BYV9(1)<br>Q9Y4A8(1) | uncertain |
| CPX-7018 | bZIP transcription factor complex, BATF-BATF3 | 2 | Q16520(1)<br>Q9NR55(1) | uncertain |
| CPX-7007 | bZIP transcription factor complex, BATF-CEBPB | 2 | P17676(1)<br>Q16520(1) | uncertain |
| CPX-7009 | bZIP transcription factor complex, BATF-CEBPD | 2 | P49716(1)<br>Q16520(1) | uncertain |
| CPX-7013 | bZIP transcription factor complex, BATF-JUND | 2 | P17535(1)<br>Q16520(1) | uncertain |
| CPX-7065 | bZIP transcription factor complex, BATF2-CEBPA | 2 | P49715(1)<br>Q8N1L9(1) | uncertain |
| CPX-7067 | bZIP transcription factor complex, BATF2-CEBPE | 2 | Q15744(1)<br>Q8N1L9(1) | uncertain |
| CPX-7066 | bZIP transcription factor complex, BATF2-CEBPG | 2 | P53567(1)<br>Q8N1L9(1) | uncertain |
| CPX-7068 | bZIP transcription factor complex, BATF2-DBP | 2 | Q10586(1)<br>Q8N1L9(1) | uncertain |
| CPX-7064 | bZIP transcription factor complex, BATF2-DDIT3 | 2 | P35638(1)<br>Q8N1L9(1) | uncertain |
| CPX-7081 | bZIP transcription factor complex, BATF2-HLF | 2 | Q16534(1)<br>Q8N1L9(1) | uncertain |
| CPX-7085 | bZIP transcription factor complex, BATF2-MAFF | 2 | Q8N1L9(1)<br>Q9ULX9(1) | uncertain |
| CPX-7096 | bZIP transcription factor complex, BATF3-CEBPB | 2 | P17676(1)<br>Q9NR55(1) | uncertain |
| CPX-7098 | bZIP transcription factor complex, BATF3-CEBPD | 2 | P49716(1)<br>Q9NR55(1) | uncertain |
| CPX-7099 | bZIP transcription factor complex, BATF3-CEBPE | 2 | Q15744(1)<br>Q9NR55(1) | uncertain |
| CPX-7109 | bZIP transcription factor complex, BATF3-CREB3 | 2 | O43889(1)<br>Q9NR55(1) | uncertain |
| CPX-7107 | bZIP transcription factor complex, BATF3-HLF | 2 | Q16534(1)<br>Q9NR55(1) | uncertain |

| | | | | |
|---|---|---|---|---|
| CPX-7103 | bZIP transcription factor complex, BATF3-MAFF | 2 | Q9NR55(1)<br>Q9ULX9(1) | uncertain |
| CPX-7105 | bZIP transcription factor complex, BATF3-MAFG | 2 | O15525(1)<br>Q9NR55(1) | uncertain |
| CPX-5342 | RXRalpha-NCOA1 activated retinoic acid receptor complex | 2 | P19793(2)<br>Q15788(2) | uncertain |

TABLE S2: **Classification of transcription factors in *H. sapiens*.** The table provides a list of 74 TFs in *H. sapiens* that belong to the basic helix-loop-helix (bHLH) or basic leucine zipper (bZIP) classes. The classes of the TFs are determined using the JASPAR database. The column 'UniProt ID' provides the UniProt IDs for each of the 74 TFs, whereas 'TF name' gives the name of the TF.

| **UniProt ID** | **TF name** | **Class** |
|---|---|---|
| Q9HBZ2 | ARNT2 | Basic helix-loop-helix factors (bHLH) |
| P50553 | ASCL1 | Basic helix-loop-helix factors (bHLH) |
| Q92858 | ATOH1 | Basic helix-loop-helix factors (bHLH) |
| Q8N100 | ATOH7 | Basic helix-loop-helix factors (bHLH) |
| O15516 | CLOCK | Basic helix-loop-helix factors (bHLH) |
| Q6QHK4 | FIGLA | Basic helix-loop-helix factors (bHLH) |
| P61296 | HAND2 | Basic helix-loop-helix factors (bHLH) |
| Q14469 | HES1 | Basic helix-loop-helix factors (bHLH) |
| Q9Y543 | HES2 | Basic helix-loop-helix factors (bHLH) |
| Q5TA89 | HES5 | Basic helix-loop-helix factors (bHLH) |
| Q96HZ4 | HES6 | Basic helix-loop-helix factors (bHLH) |
| Q9BYE0 | HES7 | Basic helix-loop-helix factors (bHLH) |
| Q9Y5J3 | HEY1 | Basic helix-loop-helix factors (bHLH) |
| Q9UBP5 | HEY2 | Basic helix-loop-helix factors (bHLH) |
| Q16665 | HIF1A | Basic helix-loop-helix factors (bHLH) |
| P61244 | MAX | Basic helix-loop-helix factors (bHLH) |
| O75030 | MITF | Basic helix-loop-helix factors (bHLH) |
| Q9UH92 | MLX | Basic helix-loop-helix factors (bHLH) |
| Q99583 | MNT | Basic helix-loop-helix factors (bHLH) |
| A6NI15 | MSGN1 | Basic helix-loop-helix factors (bHLH) |
| P50539 | MXI1 | Basic helix-loop-helix factors (bHLH) |
| P01106 | MYC | Basic helix-loop-helix factors (bHLH) |
| P04198 | MYCN | Basic helix-loop-helix factors (bHLH) |
| P13349 | MYF5 | Basic helix-loop-helix factors (bHLH) |
| P23409 | MYF6 | Basic helix-loop-helix factors (bHLH) |
| P15172 | MYOD1 | Basic helix-loop-helix factors (bHLH) |
| P15173 | MYOG | Basic helix-loop-helix factors (bHLH) |
| Q8TAK6 | OLIG1 | Basic helix-loop-helix factors (bHLH) |
| Q13516 | OLIG2 | Basic helix-loop-helix factors (bHLH) |
| Q7RTU3 | OLIG3 | Basic helix-loop-helix factors (bHLH) |
| O43680 | TCF21 | Basic helix-loop-helix factors (bHLH) |
| Q9UL49 | TCFL5 | Basic helix-loop-helix factors (bHLH) |

| | | |
|---|---|---|
| Q01664 | TFAP4 | Basic helix-loop-helix factors (bHLH) |
| P19532 | TFE3 | Basic helix-loop-helix factors (bHLH) |
| P19484 | TFEB | Basic helix-loop-helix factors (bHLH) |
| O14948 | TFEC | Basic helix-loop-helix factors (bHLH) |
| P22415 | USF1 | Basic helix-loop-helix factors (bHLH) |
| Q15853 | USF2 | Basic helix-loop-helix factors (bHLH) |
| P15336 | ATF2 | Basic leucine zipper factors (bZIP) |
| P18847 | ATF3 | Basic leucine zipper factors (bZIP) |
| P18848 | ATF4 | Basic leucine zipper factors (bZIP) |
| P17544 | ATF7 | Basic leucine zipper factors (bZIP) |
| O14867 | BACH1 | Basic leucine zipper factors (bZIP) |
| Q9BYV9 | BACH2 | Basic leucine zipper factors (bZIP) |
| Q16520 | BATF | Basic leucine zipper factors (bZIP) |
| Q9NR55 | BATF3 | Basic leucine zipper factors (bZIP) |
| P49715 | CEBPA | Basic leucine zipper factors (bZIP) |
| P17676 | CEBPB | Basic leucine zipper factors (bZIP) |
| P49716 | CEBPD | Basic leucine zipper factors (bZIP) |
| Q15744 | CEBPE | Basic leucine zipper factors (bZIP) |
| P53567 | CEBPG | Basic leucine zipper factors (bZIP) |
| P16220 | CREB1 | Basic leucine zipper factors (bZIP) |
| O43889 | CREB3 | Basic leucine zipper factors (bZIP) |
| Q03060 | CREM | Basic leucine zipper factors (bZIP) |
| Q10586 | DBP | Basic leucine zipper factors (bZIP) |
| P01100 | FOS | Basic leucine zipper factors (bZIP) |
| P15407 | FOSL1 | Basic leucine zipper factors (bZIP) |
| P15408 | FOSL2 | Basic leucine zipper factors (bZIP) |
| Q16534 | HLF | Basic leucine zipper factors (bZIP) |
| Q8WYK2 | JDP2 | Basic leucine zipper factors (bZIP) |
| P05412 | JUN | Basic leucine zipper factors (bZIP) |
| P17275 | JUNB | Basic leucine zipper factors (bZIP) |
| P17535 | JUND | Basic leucine zipper factors (bZIP) |
| O75444 | MAF | Basic leucine zipper factors (bZIP) |
| Q8NHW3 | MAFA | Basic leucine zipper factors (bZIP) |
| Q9ULX9 | MAFF | Basic leucine zipper factors (bZIP) |
| O15525 | MAFG | Basic leucine zipper factors (bZIP) |
| O60675 | MAFK | Basic leucine zipper factors (bZIP) |
| Q16621 | NFE2 | Basic leucine zipper factors (bZIP) |
| Q16649 | NFIL3 | Basic leucine zipper factors (bZIP) |
| P54845 | NRL | Basic leucine zipper factors (bZIP) |
| Q10587 | TEF | Basic leucine zipper factors (bZIP) |
| P17861 | XBP1 | Basic leucine zipper factors (bZIP) |
| Q16656 | NRF1 | Basic leucine zipper factors (bZIP) |

TABLE S3. **A list of 17 complexes from the set of 617 complexes in *S. cerevisiae* such that all the protein subunits of these complexes are transcription factors.** 'Complex ID' is the identifier of the complex as given in the EBI Complex Portal database. The 617 complexes in *S. cerevisiae* were obtained from the EBI Complex Portal database. The list of TFs was obtained from the Yeastract database. 'Complex name' is the name of the complex as given in the EBI Complex Portal database. 'Size of the complex' is the number of protein subunits the complex is constituted of. 'Uniprot ID of protein subunits' gives the Uniprot IDs of the subunits in a complex along with their stoichiometric coefficients as integers within brackets. 'Is a TR' column is 'yes' if the complex acts as a transcriptional regulator (TR) based on manual literature curation. For a given complex, if all subunits show evidence for both TR binding and effects on expression, then its entry is 'yes' in the column 'TR Binding and expression evidence', otherwise, the entry is 'no'. There are 11 complexes for which all protein subunits show evidence for both DNA binding and effects on expression.

| **Complex ID** | **Complex name** | **Size of the complex** | **Uniprot ID of protein subunits** | **Is a TR** | **TR Binding and expression evidence** |
|---|---|---|---|---|---|
| CPX-575 | Ste12/Dig1/Dig2 transcription regulation complex | 3 | P13574(0) Q03063(0) Q03373(0) | uncertain | no |
| CPX-576 | Tec1/Ste12/Dig1 transcription regulation complex | 3 | P13574(0) P18412(0) Q03063(0) | yes | no |
| CPX-828 | RTG transcription factor complex | 2 | P32607(1) P38165(1) | yes | yes |
| CPX-946 | SBF transcription complex | 2 | P09959(1) P25302(1) | yes | no |
| CPX-950 | MBP transcription complex | 2 | P09959(0) P39678(0) | yes | no |
| CPX-999 | MET4-MET28-MET31 sulfur metabolism transcription factor complex | 3 | P32389(0) P40573(0) Q03081(0) | yes | yes |
| CPX-1015 | MET4-MET28-MET32 sulfur metabolism transcription factor complex | 3 | P32389(0) P40573(0) Q12041(0) | yes | yes |
| CPX-1016 | CBF1-MET4-MET28 sulfur metabolism transcription factor complex | 3 | P17106(2) P32389(0) P40573(0) | yes | yes |
| CPX-1038 | PIP2-OAF1 transcription factor complex | 2 | P39720(1) P52960(1) | yes | yes |
| CPX-1042 | GAL3-GAL80 transcription regulation complex | 2 | P04387(2) P13045(2) | yes | no |
| CPX-1044 | GAL4-GAL80 transcription repressor complex | 2 | P04386(2) P04387(2) | yes | yes |
| CPX-1200 | RAP1-GCR1 transcription activation complex | 2 | P07261(2) P11938(0) | yes | yes |
| CPX-1229 | RAP1-GCR1-GCR2 transcription activation complex | 3 | P07261(2) P11938(0) Q01722(2) | yes | yes |
| CPX-1277 | INO2-INO4 transcription activation complex | 2 | P13902(0) P26798(0) | yes | yes |
| CPX-1415 | IME1-UME6 transcription activation complex | 2 | P21190(0) P39001(0) | yes | yes |
| CPX-1663 | CYP8-TUP1 corepressor complex | 2 | P14922(1) P16649(4) | corepressor | no |
| CPX-1830 | CCAAT-binding factor complex | 4 | P06774(1) P13434(1) P14064(1) Q02516(1) | yes | yes |

TABLE S4. **Classification of transcription factors in *S. cerevisiae*** The table provides a list of 17 TFs from the Yeastract database, that belong to the basic helix-loop-helix (bHLH) or basic leucine zipper (bZIP) classes. The classes of the TFs are determined using the JASPAR database. The column 'UniProt ID' provides the UniProt IDs for each of the 17 TFs, whereas 'TF name' corresponds to the name of the TF. All the 17 TFs shown in the table display evidence for DNA binding as well as effects on expression.

| UniProt ID | TF Name | DNA binding and expression evidence | Class |
|---|---|---|---|
| P33122 | TYE7 | Yes | Basic helix-loop-helix factors (bHLH) |
| P38165 | RTG3 | Yes | Basic helix-loop-helix factors (bHLH) |
| P13902 | INO4 | Yes | Basic helix-loop-helix factors (bHLH) |
| P26798 | INO2 | Yes | Basic helix-loop-helix factors (bHLH) |
| P07270 | PHO4 | Yes | Basic helix-loop-helix factors (bHLH) |
| P17106 | CBF1 | Yes | Basic helix-loop-helix factors (bHLH) |
| P14164 | ABF1 | Yes | Basic helix-loop-helix factors (bHLH) |
| P32389 | MET4 | Yes | Basic leucine zipper factors (bZIP) |
| P40573 | MET28 | Yes | Basic leucine zipper factors (bZIP) |
| P41546 | HAC1 | Yes | Basic leucine zipper factors (bZIP) |
| P03069 | GCN4 | Yes | Basic leucine zipper factors (bZIP) |
| Q02100 | SKO1 | Yes | Basic leucine zipper factors (bZIP) |
| Q06596 | ARR1 | Yes | Basic leucine zipper factors (bZIP) |
| Q08182 | YAP7 | Yes | Basic leucine zipper factors (bZIP) |
| Q03935 | YAP6 | Yes | Basic leucine zipper factors (bZIP) |
| P40574 | YAP5 | Yes | Basic leucine zipper factors (bZIP) |
| P38749 | YAP3 | Yes | Basic leucine zipper factors (bZIP) |

TABLE S5. **Comparison of the fractions of four types of biologically meaningful BFs and the fraction of the BFs allowed by the most restrictive composition structures for $k \leq 5$ inputs.** The fractions are computed with respect to all possible BFs for $k \leq 5$ inputs. The four types of biologically meaningful BFs include Unate functions (UF), Canalyzing functions (CF), Nested canalyzing functions (NCF) and Read-once functions (RoFs). The column 'Composed BF' represents BFs contained in the most restrictive composition structure for a given $k$. The most restrictive composition structure is the composition structure that has the least number of BFs when compared to other composition structures with the same number of inputs $k$. For $k = 1$ and 2, there are no restrictions in possible BFs due to composition structure since only trivial composition structures such as $\{1\}$, $\{1,1\}$ and $\{2\}$ exist. For $k = 3$, 4 and 5, the most restrictive composition structures are $\{1,2\}$, $\{2,2\}$ and $\{2,3\}$, respectively.

| Inputs ($k$) | Fraction of | | | | |
|---|---|---|---|---|---|
| | UF | CF | NCF | RoF | Composed BF |
| 1 | 1 | 1 | 0.5 | 0.5 | 1 |
| 2 | 0.875 | 0.875 | 0.5 | 0.5 | 1 |
| 3 | 0.406 | 0.469 | 0.25 | 0.25 | 0.594 |
| 4 | 0.033 | 0.054 | 0.011 | 0.013 | 0.018 |
| 5 | $5.37 \times 10^{-5}$ | $3.01 \times 10^{-4}$ | $2.47 \times 10^{-6}$ | $3.52 \times 10^{-6}$ | $1.67 \times 10^{-5}$ |

TABLE S6. **Fraction of BFs in different composition structures that display biologically meaningful properties.** The fraction of BFs in different non-trivial composition structures that also belong to each of the four types of biologically meaningful BFs, namely Unate functions (UF), Canalyzing functions (CF), Nested canalyzing functions (NCF) and Read-once functions (RoFs). The fraction is computed with respect to the allowed BFs in a composition structure.

| **Composition structure** | **Fraction of biologically meaningful BFs in composition structure** | | | |
|---|---|---|---|---|
| | **UF** | **CF** | **NCF** | **RoF** |
| {1,2} | 0.632 | 0.789 | 0.421 | 0.421 |
| {1,3} | 0.249 | 0.722 | 0.151 | 0.151 |
| {2,2} | 0.525 | 0.604 | 0.185 | 0.265 |
| {1,1,2} | 0.220 | 0.298 | 0.118 | 0.134 |
| {1,4} | 0.022 | 0.672 | 0.006 | 0.007 |
| {2,3} | 0.191 | 0.469 | 0.046 | 0.095 |
| {1,1,3} | 0.099 | 0.307 | 0.040 | 0.054 |
| {1,2,2} | 0.177 | 0.250 | 0.055 | 0.099 |
| {1,1,1,2} | 0.018 | 0.036 | 0.003 | 0.004 |

TABLE S7. **Number and fraction of BFs with odd bias in different composition structures.** The fractions of BFs with odd bias in a composition structure are computed with respect to all allowed BFs in the composition structure. The column 'Number of composed BFs' gives the number of allowed BFs in a composition structure. The column 'Odd biases present' gives the list of odd biases of BFs that are present in a composition structure. Note that the table only gives data for non-trivial composition structures with $k \leq 5$ inputs.

| **Composition structure** | **Number of composed BFs** | **BFs with odd bias in composition structure** | | |
|---|---|---|---|---|
| | | **Number** | **Fraction** | **Odd biases present** |
| {1,2} | 152 | 64 | 0.421 | 1,3 |
| {1,3} | 4864 | 1760 | 0.361 | 1,3,5,7 |
| {2,2} | 1208 | 320 | 0.264 | 1,3,7 |
| {1,1,2} | 6216 | 2368 | 0.381 | 1,3,5,7 |
| {1,4} | 1921928 | 646144 | 0.336 | 1,3,5,7,9,11,13,15 |
| {2,3} | 71608 | 17024 | 0.238 | 1,3,5,7,9,11,15 |
| {1,1,3} | 263488 | 75584 | 0.287 | 1,3,5,7,9,11,13,15 |
| {1,2,2} | 100768 | 25344 | 0.252 | 1,3,5,7,9,11,13,15 |
| {1,1,1,2} | 3446488 | 1266944 | 0.368 | 1,3,5,7,9,11,13,15 |

TABLE S8. **Enrichment of composed BFs in the reference biological dataset.** The enrichment factors for composed BFs in different non-trivial composition structures with number of inputs $k \leq 5$ and the associated one-sided $p$-values.

| Composition structure | Enrichment factor | $p$-value |
|---|---|---|
| {1,2} | 1.63 | $6.15 \times 10^{-72}$ |
| {1,3} | 12.48 | $4.33 \times 10^{-245}$ |
| {2,2} | 40.37 | $3.90 \times 10^{-274}$ |
| {1,1,2} | 10.30 | $7.57 \times 10^{-250}$ |
| {1,4} | 1948.23 | 0 |
| {2,3} | 45760.08 | 0 |
| {1,1,3} | 14732.62 | 0 |
| {1,2,2} | 36887.66 | 0 |
| {1,1,1,2} | 1158.31 | 0 |

TABLE S9. **$p$-values corresponding to the relative enrichment values of biologically meaningful BFs within different composition structures.** The $p$-values corresponding to the relative enrichment values $E_R$ for the four biologically meaningful sub-types of BFs within different composition structures with number of inputs $k \leq 5$. The four biologically meaningful sub-types within composed BFs include those BFs in a composition structure that also happen to be Unate functions (UF), Canalyzing functions (CF), Nested canalyzing functions (NCF), or Read-once functions (RoFs).

| Composition structure | $p$-values corresponding to $E_R$ of sub-types in composition structure | | | |
|---|---|---|---|---|
| | UF | CF | NCF | RoF |
| {1,2} | 0 | 0 | $1.79 \times 10^{-115}$ | $1.79 \times 10^{-115}$ |
| {1,3} | 0 | 0 | $2.27 \times 10^{-176}$ | $2.27 \times 10^{-176}$ |
| {2,2} | $1.75 \times 10^{-54}$ | $1.98 \times 10^{-24}$ | $5.61 \times 10^{-100}$ | $3.74 \times 10^{-96}$ |
| {1,1,2} | $3.08 \times 10^{-166}$ | $1.30 \times 10^{-105}$ | $1.42 \times 10^{-185}$ | $4.53 \times 10^{-202}$ |
| {1,4} | $5.19 \times 10^{-227}$ | 0 | $2.26 \times 10^{-254}$ | $3.13 \times 10^{-258}$ |
| {2,3} | 0 | $1.74 \times 10^{-27}$ | $1.62 \times 10^{-111}$ | $6.60 \times 10^{-105}$ |
| {1,1,3} | 0 | $1.26 \times 10^{-57}$ | $7.92 \times 10^{-146}$ | $8.03 \times 10^{-160}$ |
| {1,2,2} | 0 | $4.12 \times 10^{-64}$ | $7.77 \times 10^{-123}$ | $2.05 \times 10^{-118}$ |
| {1,1,1,2} | 0 | $1.32 \times 10^{-181}$ | $2.18 \times 10^{-277}$ | $9.30 \times 10^{-301}$ |

TABLE S10. **$p$-values for comparison between the enrichments of composed BFs and biologically meaningful BFs with minimum complexity in the reference biological dataset.** $T_C$ denotes the set of composed BFs allowed by a composition structure at a given number of inputs $k$, $T_{NCF}$ denotes the set of all $k$-input nested canalyzing functions (NCFs), and $T_{RoF}$ denotes the set of all $k$-input Read-once functions (RoFs). $\cap$ represents the intersection of two sets and $\setminus$ represents the set-theoretic difference. "–" in the columns $T_{NCF} \setminus T_C$ or $T_{RoF} \setminus T_C$ indicates that the NCFs or RoFs are a subset of the set of BFs allowed by the composition structure.

| **Composition structure** | $T_C \cap T_{NCF}$ | $T_C \setminus T_{NCF}$ | $T_{NCF} \setminus T_C$ | $T_C \cap T_{RoF}$ | $T_C \setminus T_{RoF}$ | $T_{RoF} \setminus T_C$ |
|---|---|---|---|---|---|---|
| {1,2} | $3.09 \times 10^{-184}$ | 1 | – | $3.09 \times 10^{-184}$ | 1 | – |
| {1,3} | 0 | 0.966 | – | 0 | 0.966 | $1.66 \times 10^{-19}$ |
| {2,2} | 0 | $1.59 \times 10^{-11}$ | $7.07 \times 10^{-70}$ | 0 | 0.009 | $7.07 \times 10^{-70}$ |
| {1,1,2} | 0 | 0.406 | – | 0 | 0.999 | – |
| {1,4} | 0 | $2.17 \times 10^{-35}$ | – | 0 | $7.77 \times 10^{-26}$ | $1.06 \times 10^{-47}$ |
| {2,3} | 0 | $9.10 \times 10^{-80}$ | $4.44 \times 10^{-106}$ | 0 | $4.83 \times 10^{-30}$ | $8.09 \times 10^{-105}$ |
| {1,1,3} | 0 | $2.87 \times 10^{-67}$ | – | 0 | $8.85 \times 10^{-25}$ | $3.43 \times 10^{-05}$ |
| {1,2,2} | 0 | $1.36 \times 10^{-76}$ | $1.31 \times 10^{-30}$ | 0 | $1.00 \times 10^{-28}$ | $1.31 \times 10^{-30}$ |
| {1,1,1,2} | 0 | $7.82 \times 10^{-52}$ | – | 0 | $1.51 \times 10^{-22}$ | – |

TABLE S11. **Comparison between the enrichments of composed BFs and biologically meaningful BFs in the reference biological dataset.** The table provides the enrichment factors when composed BFs allowed by non-trivial composition structures with $k \leq 5$ inputs are compared with two biologically meaningful BFs namely unate functions (UFs) and canalyzing functions (CFs). $T_C$ denotes the set of composed BFs allowed by a composition structure at a given number of inputs $k$, whereas $T_{UF}$ and $T_{CF}$ denote the set of all $k$-input UFs and CFs, respectively. $\cap$ represents the intersection of two sets and $\setminus$ represents the set-theoretic difference. "–" in the column $T_{CF} \setminus T_C$ indicates that the CFs are a subset of the set of BFs allowed by the corresponding composition structure.

| **Composition structure** | $T_C \cap T_{UF}$ | $T_C \setminus T_{UF}$ | $T_{UF} \setminus T_C$ | $T_C \cap T_{CF}$ | $T_C \setminus T_{CF}$ | $T_{CF} \setminus T_C$ |
|---|---|---|---|---|---|---|
| {1,2} | 2.58 | 0.00 | 1.01 | 2.06 | 0.00 | – |
| {1,3} | 50.17 | 0.00 | 4.76 | 17.28 | 0.00 | – |
| {2,2} | 76.53 | 0.44 | 10.91 | 61.59 | 7.97 | 5.66 |
| {1,1,2} | 46.54 | 0.05 | 1.91 | 32.54 | 0.87 | 0.31 |
| {1,4} | 89183.19 | 14.64 | 2623.97 | 2897.47 | 0.00 | – |
| {2,3} | 239564.87 | 0.00 | 4316.45 | 90144.74 | 6518.64 | 568.71 |
| {1,1,3} | 148416.79 | 0.00 | 1616.48 | 44868.92 | 1357.79 | 90.92 |
| {1,2,2} | 208387.45 | 0.00 | 2329.87 | 136371.86 | 3645.06 | 239.02 |
| {1,1,1,2} | 64895.59 | 0.00 | 1303.10 | 30112.53 | 91.11 | 47.07 |

TABLE S12. $p$**-values for comparison between the enrichments of composed BFs and biologically meaningful BFs in the reference biological dataset.** The table provides the enrichment factors when composed BFs allowed by non-trivial composition structures with $k \leq 5$ inputs are compared with two biologically meaningful BFs namely, unate functions (UFs) and canalyzing functions (CFs). $T_C$ denotes the set of composed BFs allowed by a composition structure at a given number of inputs $k$, whereas $T_{UF}$ and $T_{CF}$ denote the set of all $k$-input UFs and CFs, respectively. $\cap$ represents the intersection of two sets and $\setminus$ represents the set-theoretic difference. "–" in the column $T_{CF} \setminus T_C$ indicates that the CFs are a subset of the set of BFs allowed by the corresponding composition structure.

| **Composition structure** | $T_C \cap T_{UF}$ | $T_C \setminus T_{UF}$ | $T_{UF} \setminus T_C$ | $T_C \cap T_{CF}$ | $T_C \setminus T_{CF}$ | $T_{CF} \setminus T_C$ |
|---|---|---|---|---|---|---|
| {1,2} | $3.23 \times 10^{-149}$ | 1 | 0.413 | $1.83 \times 10^{-111}$ | 1 | – |
| {1,3} | 0 | 0.999 | $1.39 \times 10^{-08}$ | $8.86 \times 10^{-279}$ | 0.995 | – |
| {2,2} | 0 | 0.661 | $5.93 \times 10^{-49}$ | $1.14 \times 10^{-280}$ | $1.39 \times 10^{-10}$ | $1.02 \times 10^{-29}$ |
| {1,1,2} | 0 | 0.999 | 0.041 | 0 | 0.652 | 0.960 |
| {1,4} | 0 | 0.002 | $2.02 \times 10^{-59}$ | 0 | 0.023 | – |
| {2,3} | 0 | 0.002 | $3.64 \times 10^{-116}$ | 0 | $5.16 \times 10^{-36}$ | $5.39 \times 10^{-66}$ |
| {1,1,3} | 0 | 0.009 | $1.99 \times 10^{-38}$ | 0 | $3.34 \times 10^{-29}$ | $1.24 \times 10^{-09}$ |
| {1,2,2} | 0 | 0.003 | $1.93 \times 10^{-58}$ | 0 | $1.16 \times 10^{-36}$ | $1.17 \times 10^{-25}$ |
| {1,1,1,2} | 0 | 0.116 | $2.68 \times 10^{-26}$ | 0 | $1.17 \times 10^{-20}$ | $1.22 \times 10^{-05}$ |
